# p53 rapidly restructures 3D chromatin organization to trigger a transcriptional response

**DOI:** 10.1101/2023.10.10.561663

**Authors:** François Serra, Andrea Nieto-Aliseda, Lucía Fanlo, Llorenç Rovirosa, Mónica Cabrera-Pasadas, Aleksey Lazarenkov, Blanca Urmeneta, Álvaro Alcalde, Emanuele M. Nola, Andrei L. Okorokov, Peter Fraser, Mariona Graupera, Sandra D. Castillo, Jose Luis Sardina, Alfonso Valencia, Biola M. Javierre

**Affiliations:** Josep Carreras Leukaemia Research Institute, Badalona, Barcelona, Spain; Barcelona Supercomputing Center, Barcelona, Barcelona, Spain; Wolfson Institute for Biomedical Research, University College London, London, UK; Department of Biological Science, Florida State University, Tallahassee, FL, USA; Catalan Institution for Research and Advanced Studies (ICREA), Barcelona, Spain; CIBERONC, Instituto de Salud Carlos III, Madrid, Spain; Institute for Health Science Research Germans Trias i Pujol, Badalona, Barcelona, Spain

## Abstract

Activation of the p53 tumor suppressor triggers a transcriptional program to control cellular response to stress. However, the molecular mechanisms by which p53 controls gene transcription are not completely understood. Here, using a multi-omics integration framework, we uncover the critical role of spatio-temporal genome architecture in this process. We demonstrate that p53 drives direct and indirect changes in genome compartments, topologically associating domains and DNA loops within minutes of its activation, which escort the p53 transcriptional program along time. Focused on p53-bound enhancers, we report a core transcriptional program of 340 genes directly regulated by p53 over distance, most of these not previously identified. Finally, we showcase that p53 controls transcription of distal genes through newly formed and pre-existing enhancer-promoter loops in a cohesin dependent manner. Taken together, our findings demonstrate a previously unappreciated architectural role of p53 as regulator at distinct topological layers and provide a reliable set of new p53 direct target genes that may help future designs of p53-based cancer therapies.

---

The *TP53* tumor suppressor gene, colloquially known as the “guardian of the genome”, is the most frequently altered gene in cancer. It encodes for a sequence-specific transcription factor named p53 that, after its activations via cell stress or DNA damage, triggers transcriptional activation of a myriad of target genes to ultimately facilitate distinct cellular outcomes, including cell cycle arrest, senescence or apoptosis among others^1^. However, the molecular mechanisms by which p53 controls gene transcription are not completely understood despite being the most studied gene in history.

Gene transcription is intimately associated with the three-dimensional (3D) packing of the chromatin within the nucleus and its alteration is closely linked to disease^2^. Chromatin folding involves three hierarchical levels. First, at megabases scale, the genome can be segregated into the so-called A and B compartments. The A compartment represents active, accessible chromatin with a tendency to occupy a more central position in the nucleus. The B compartment corresponds to heterochromatin and is enriched at the nuclear periphery. Second, topologically associating domains (TADs) are sub-megabase structures delimitated by TAD borders that interact more frequently within themselves than with the rest of the genome. TADs are conserved across species and cell types and show a coordinated transcriptional status. Third, these domains are formed by assemblies of chromatin loops^3^. Among hierarchical levels, chromatin loops engaging gene promoters and distal regulatory elements (*e.g.,* enhancers or silencers) are key to control gene transcription since these often locate distally in the genome but need physical proximity to function^4^. Despite its critical role in gene transcription regulation, the interplay between spatio-temporal genome organization and p53 remains largely unexplored. Besides, p53 frequently binds enhancers and most of its linked distal target genes are unknown. Study of the spatio-temporal genome organization represents thus a direct way to identify the true set of genes directly regulated by p53.

Transcription factor CCCTC-binding factor (CTCF) and the ring-shaped multiprotein complex cohesin, which includes the major subunit RAD21, play key roles in establishing TADs and loops thought a process called ATP-dependent loop extrusion. In this process, the cohesin complex is loaded onto DNA and pumps it until encountering a barrier, which is often formed by CTCF^5–9^. Therefore, cohesin degradation leads to rapid disappearance of TADs, antagonizes compartment segregation and promotes loss of most of the significant promoter-anchored interactions^6,9–17^. Surprisingly, complete removal of cohesin has little impact on steady-state gene transcription^6^ but prevents adequate activation of inflammatory gene response^18^ and initiation of new regulatory programs in neurons acquiring response to new stimuli^19^. However, the role of cohesin in activation of other types of inducible genes, such as those activated by p53, remains largely unexplored^20–22^.

Here, we addressed for the first time the relationship between p53 activation and spatio- temporal genome architecture to establish a transcriptional response to cellular stress. First, we uncovered a new role of p53 as a master remodeler of the spatio-temporal chromatin organization. Unexpectedly, p53-driven topological changes, including the formation of long- range DNA loops between p53-bound enhancers and promoters, occur minutes after its activation and enable the establishment of the p53 transcriptional response. Second, we discovered an unforeseen dependence of p53 inducible gene expression on cohesin. Finally, we identified a new set of direct target genes transcriptionally controlled by p53 over distance. Taken together, our results demonstrate the power of integrating 3D genome architecture, epigenetics, gene expression and transcription factor binding profiles along time to gain insight into gene regulatory mechanisms and to discover new transcription factor target genes.

## Results

### p53 activation drives two waves of dramatic changes in genome compartments and TADs

To address global consequences of p53 activation in 3D genome organization we used the HCT116 cell line, a widely used model characterized by a wild type p53 response. These cells were treated with 10µM Nutlin-3a, an inhibitor of the p53-HDM2 interaction used to mimic cell stress-induced p53 activation, and collected at five time points (1, 4, 7, 10 and 24 hours post treatment). We also treated cells with the drug vehicle (*i.e.,* DMSO) and used these cells as a control of basal conditions without p53 activation (also referred to in the manuscript as 0 hours of Nutlin-3a) (Fig. 1A and Supplementary Fig. 1 A-C). We used *in situ* Hi-C to generate genome-wide chromosome conformation maps along all six time points of p53 activation (Supplementary Data 1). After confirming libraries quality and reproducibility between biological replicates (SCC > 0.95; see methods) (Fig. 1B and Supplementary Fig. 1D), we segmented the genome into A and B compartments (Supplementary Fig. 1E). Clustering of biological replicates and separation of treatment conditions demonstrated robust and dynamic A/B compartmentalization associated with p53 activation (Fig. 1C and Supplementary Fig. 1F- G). Globally, although most of the genome compartments remained stable throughout all time points, 12.5% of A or B compartments switched during p53 activation, from here on out referred to as dynamic compartments (Fig. 1D). Furthermore, per time points, we observed compartment activation at 1 hour, limited compartment rewiring from 1 to 7 hours and compartment inactivation at 10 hours of p53 activation onwards (Fig. 1E-H and Supplementary Fig. 1H). Specifically, clustering based on compartment score identified 7 main clusters of dynamic compartments following p53 activation. Interestingly, 23.2% of these (cluster 1 and 2) dramatically flipped from B to A after 1 hour of Nutlin-3a treatment and then progressively reduced their compartment score switching back to B. Contrarily, 39.5% of the dynamic compartments (clusters 3, 4 and 5) were A compartments specifically inactivated at late time points (10 or 24 hours of Nutlin-3a treatment). However, these also increased their compartment score after 1 hour of p53 activation despite already being classified as A compartments before p53 activation (*i.e.*, 0 hours), reinforcing the observed unexpected tendency of early, genome-wide gain of genome activity.

**Fig 1.**
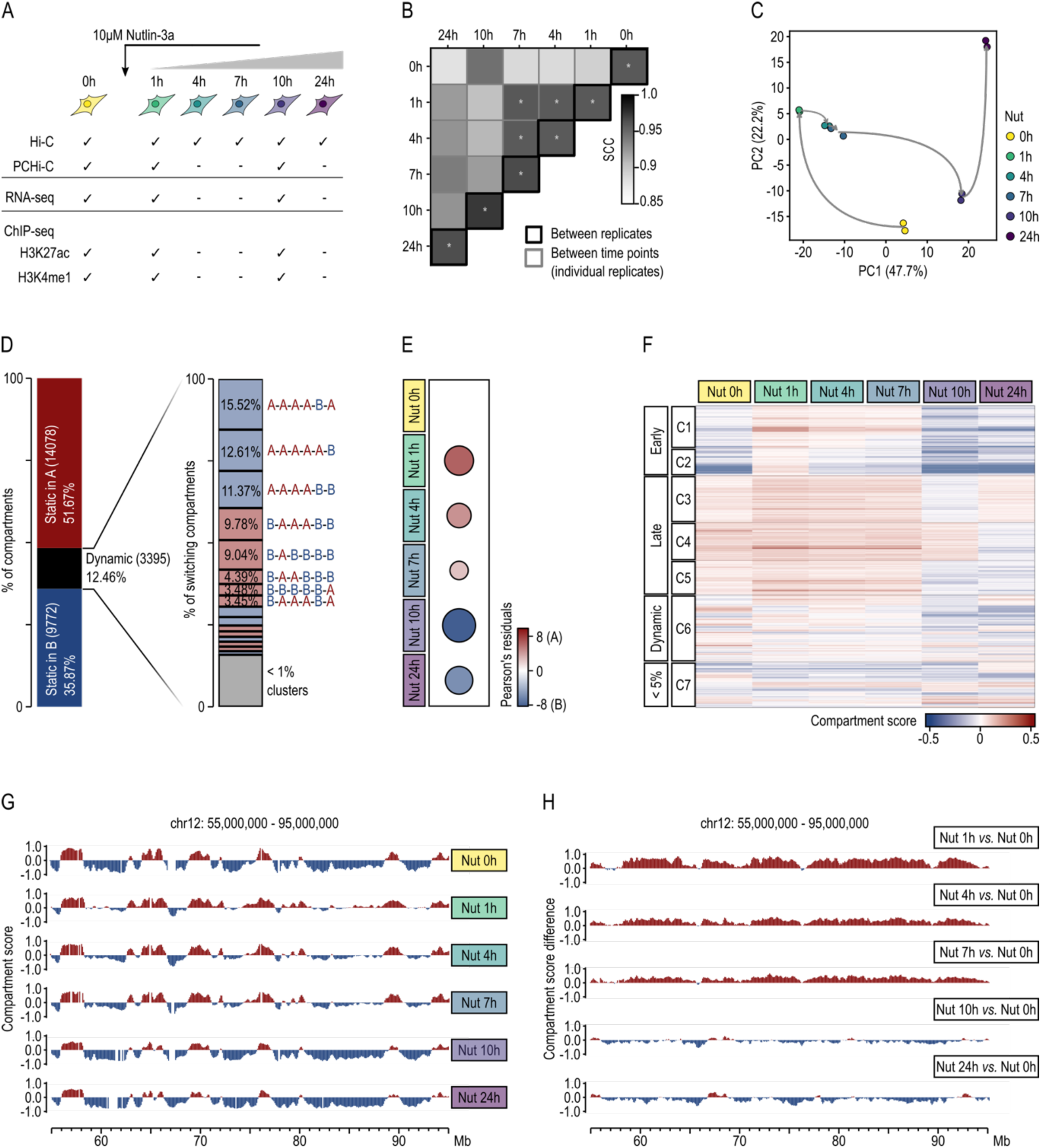
A/B compartment changes along p53 activation. **A.** Experimental multi-omics framework of the work. 0h means control cells treated with drug vehicle characterized by no p53 activation. **B.** Similarity between Hi-C interaction matrices calculated using the Stratum Adjusted Correlation Coefficient (SCC) across the time course. SCC value of 1 means perfect correlation, values above 0.95 (represented by a star) are expected for biological replicates. **C.** Principal Component Analysis of the compartment scores for each Hi-C individual biological replicate across p53 activation. Principal components 1 and 2 were taken into consideration. Numbers in parentheses represent the percentage of variance explained by each principal component (PC). **D.** Percentage of compartment categories according to their dynamics along p53 activation. Dynamic compartments are defined as those that change compartment between any two time points. **E.** Correlation plot for resultant Pearson’s residuals from a Chi-square test of independence for the contingency table time-course *vs.* compartment category (X-squared = 410.52, df = 5, p- value < 2.2e-16). Only significant associations are shown, where significance is determined as |Pearson’s residual| > 2. Bubble size indicates the number of compartment regions associated, and blue to red corresponds to an increasing association with A compartments and a decreasing association with B compartments. **F.** Compartment scores of the dynamic compartments in response to p53 activation, with each row representing a 100kb bin and each column representing a time point. The heatmap color scale represents the compartment score assigned to each 100kb bin with positive values indicating A compartment and negative values indicating B compartment. **G.** Variation of eigenvector values along a 40 Mb genomic region of chromosome 12. The genomic bins are colored according to their compartment type. **H.** Difference in compartment score between p53 activated samples and non-activated cells (Nut 0h) along the genomic regions displayed on panel G. Positive values are represented in red and indicate compartment activation (*i.e.,* shift towards the A compartment) along p53 activation, while negative values are in blue and indicate a shift towards the B compartment.

Next, we used chromosome-wide insulation potential (TADbit score > 4^23^) to identify between 2,820 and 3,963 TADs per time point with a median size ranging from 453 to 622 kilobases (Fig. 2A-B). Genome-wide insulation scores analyzed by Principal Component Analysis (PCA) and hierarchical clustering over time revealed high reproducibility between biological replicates and progressive changes reflecting p53 activation (Fig. 1C and Supplementary Fig. 1I-J). While 2,091 of TAD borders were stable across all stages, the remaining ones were dynamically lost or gained. Similar to A/B compartments, 1 hour of p53 activation promoted a dramatic architectural change, this time resulting in loss of TAD borders (Fig. 2D-E and Supplementary Fig. 1K). TAD profiles were then extensively conserved until the second wave of rewiring occurring after 10 hours of treatment. At this stage, we observed a sharp acquisition of TAD borders followed by, after 24 hours of treatment, a global erase of these topological features. Specifically, 26.3% of the dynamic TAD borders rapidly disappeared after 1 hour of treatment (clusters 2 and 4, Fig. 2E). One third of these (cluster 4) temporally reappeared after 10 hours of p53 activation before disappearing again at 24h. Moreover, 16.9% and 8.1% of the dynamic TAD borders specifically disappeared (cluster 3 and 4) or appeared (cluster 5) at 24 hours of treatment, respectively. Besides these trends of appearance and disappearance, we observed a global and progressive increase of the insulation capacity of TAD borders - according to their insulation scores^24^ -, suggesting that less insulating borders are preferentially erased during p53 activation (Fig. 2F).

**Fig 2.**
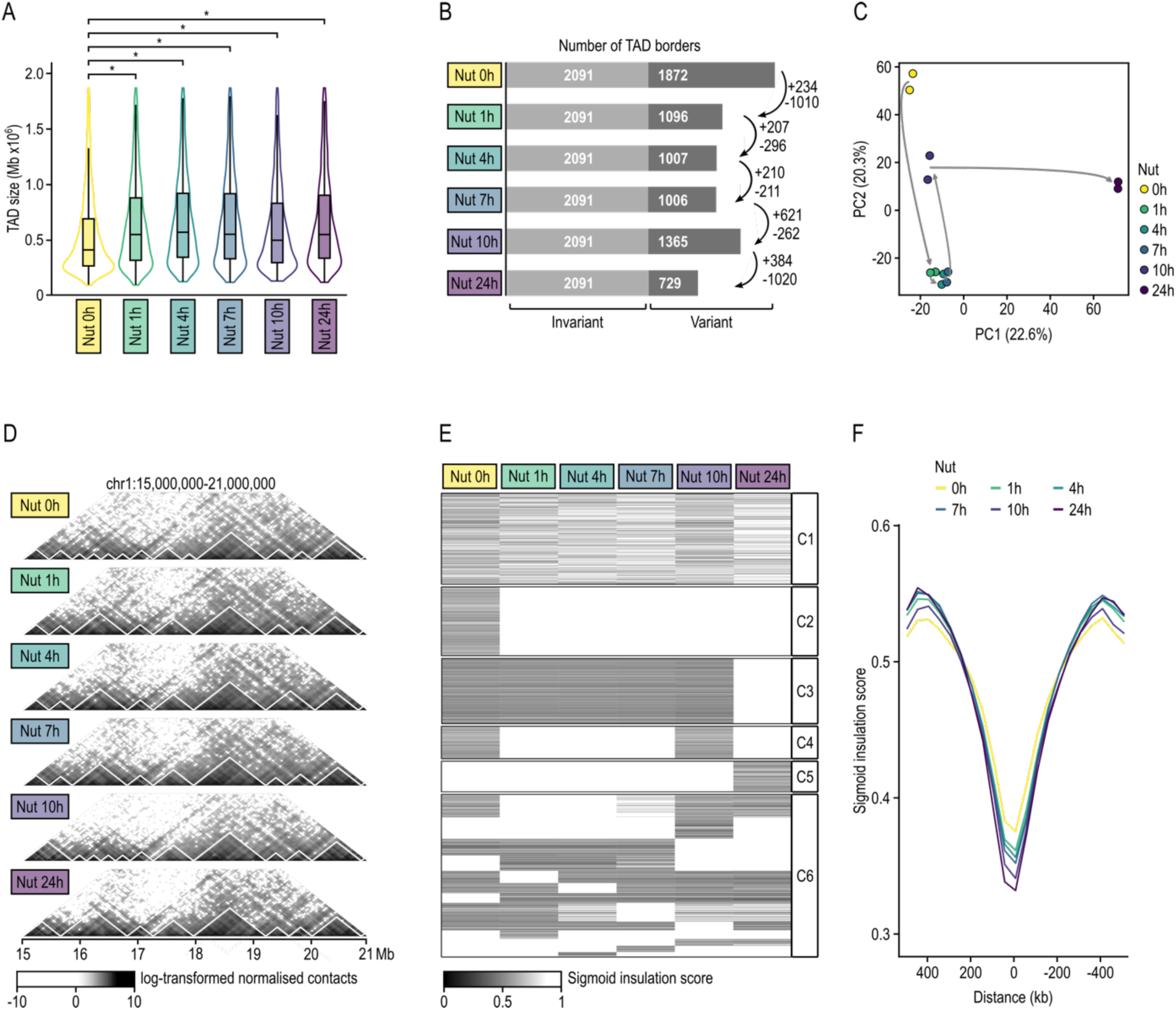
TAD dynamics along p53 activation. **A.** Distribution of TAD sizes along p53 activation. A two-sided Wilcoxon test was used to test whether distributions of TAD sizes differed throughout p53 activation compared to control. Nut 0h refers to control cells without p53 activation. **B.** Bar plot displaying the number of variant and invariant TAD borders along p53 activation. Invariant TAD borders are those detected at all time points. Positive/negative numbers near arrows represent the number of TAD borders gained/lost between the corresponding time points. **C.** Principal Component Analysis of TADbit TAD border scores for all Hi-C biological replicates across p53 activation. Principal components 1 and 2 were taken into consideration. Numbers in parentheses represent the percentage of variance explained by each principal component (PC). **D.** Hi-C contact maps of a 6Mb genomic region of chromosome 1 where TAD are marked by white lines. **E.** Clustering of TAD borders based on their TAD insulation scores and the patterns of temporal changes observed throughout p53 activation. Only TAD borders characterized by a TADbit score > 4 were included. TAD borders were manually clustered considering their dynamism throughout p53 activation. **F.** Insulation score profiles stacked over a 1Mb genomic region centered at TAD borders. Only TAD borders characterized by a TADbit score > 4 were included. Sigmoid insulation score is inversely proportional to insulation capacity.

Altogether, these results demonstrate dramatic changes in genome architecture during p53 activation, including early changes at 1 hour of activation and late changes after 10 hours of treatment, which may be direct or indirect downstream effects of p53 pathway’s activation.

### Early and late genome organization changes are triggered by p53 binding directly and indirectly

Since p53 activation leads to two waves of genome organization dynamics, we decided to explore the underlying molecular mechanisms driving these sharp transitions. To do so, using cells collected after 1 and 10 hours of Nutlin-3a treatment and control cells treated with the drug vehicle (0 hours), we generated: i) high quality chromatin immunoprecipitation with massively parallel sequencing (ChIP-seq) libraries of two histone modifications (H3K4me1 and H3K27ac), enabling the identification of primed enhancers (H3K4me1), active enhancers (H3K4me1 and H3K27ac) and active promoters (H3K27ac) (Supplementary Data 2); and ii) RNA-seq libraries that enable the genome-wide profiling of gene transcription (Fig. 1A and Supplementary Data 3). Besides, we also used already processed publicly available ChIP-seq data of p53’s binding profile in the same cell type subjected to similar activation^25^.

After verifying libraries’ quality and reproducibility between biological replicates (Supplementary Fig. 2A-K), we identified enhancers and defined their activities along p53 activation (Supplementary Fig. 3A-D and Supplementary Data 4). Next, we tested whether changes induced by Nutlin-3a treatment were direct (*i.e.,* local shifts provoked by p53 binding) or indirect effects of p53 (*e.g.,* alterations through transcription factors regulated by p53). p53 binding to chromatin is largely invariant but not all binding events deliver transactivation, leading to cell type and stimulus-specific variation in the p53 transcriptional program^26–28^. Thus, we defined as functional p53 binding sites a set of 2,105 sites bound by p53 and characterized by the presence of the activating H3K27ac histone mark in the time points corresponding to p53 activation condition (*i.e.,* 1 or 10 hours of Nutlin-3a treatment) (Supplementary Fig. 3E-G and Supplementary Data 5). Only 6.5% (137/2,105) of those functional p53 bindings occurred at promoters (*i.e.,* -1000 +200bp of any TSS). Genes involved in these bindings had some degree of activity at basal conditions (*i.e.,* H3K27ac at promoters), and tended to gain promoter H3K27ac deposition and gene transcription levels after Nutlin-3a treatment (Supplementary Fig. 3G-I). The remaining functional p53 binding events were distal from TSS, and among them, 901 occurred at active enhancers under p53 activation condition (*i.e.,* co-presence of H3K4me1 and H3K27ac at 1 or 10 hours of Nutlin-3a treatment). Interestingly, most of these active enhancers were already established at basal conditions and increased their H3K27ac levels as consequence of p53 activation (Supplementary Fig. 3G, J). Collectively, these results highlight p53’s preference to bind active regulatory elements and suggest its limited capacity of epigenetic remodeling on the linear genome.

Since p53 preferentially acts on pre-established regulatory elements, we explored whether its functional binding may promote topological perturbations to ultimately explain the transcriptional response. Globally, functional p53 preferentially bound A compartments (Fig. 3A) and TAD border vicinities (Fig. 3B). Interestingly, its binding was mainly enriched in compartments that gained activity during p53 activation (Fig. 3C). Indeed, compartments switching from B to A show a significant enrichment in functional p53 binding sites with respect to those switching from A to B. Indeed, after 1 hour and 10 hours of activation, 5.7% (61 out of 1,078; Fisher O.R. 3.5 p-value 0.007) and 13.1% (22 out of 168; Fisher OR 3.9 p-value 4e-6) of the B to A switched compartments, respectively, were characterized by harboring functional p53 binding sites (pink area in Fig. 3D-E). Besides, regions harboring functional p53 binding sites – independently of the compartment dynamics – tended to suffer a higher increase in compartment score (Fig. 3F-G), a higher gain of activating H3K27ac mark (Fig. 3H), and increased transcription (Fig. 3I) compared to their counterparts depleted of functional p53 bindings. Contrarily, only 1.7% (7 out of 411) and 3.7% (48 out of 1281) of the compartments that switched from A to B after 1 or 10 hours of activation, respectively, harbor functional p53 binding sites, although these tended to have a more moderate decrease in compartment score than compartments depleted of functional p53 binding (light blue area in Fig. 3E, G).

**Fig 3.**
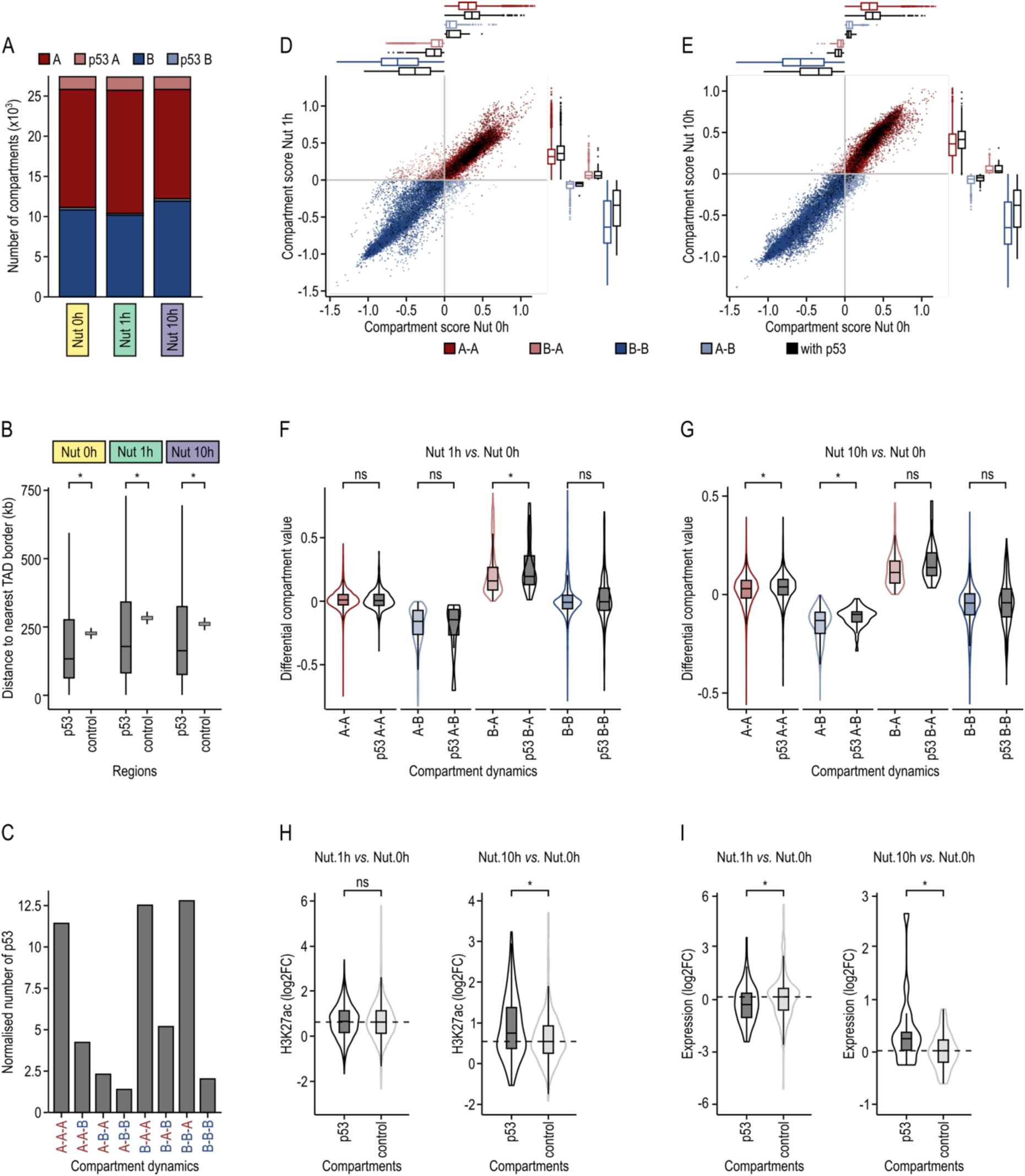
p53 binding to chromatin leads to direct changes in 3D genome topology. **A.** Proportion of functional p53 bindings in A/B compartments along p53 activation. p53A/p53B are 100kb bins in A/B compartment with at least one p53 functional binding. Nut 0h refers to control cells without p53 activation. **B.** Distribution of the distances between functional “p53” binding sites, or “control” regions, and the nearest TAD border. Control regions are defined as a random set of regions of similar genomic length to a p53 binding site and a similar distribution across the genome. Statistical comparisons between p53 and control regions were computed using a two-sided Wilcoxon test. **C.** Proportion of p53 bindings in the different categories of A/B compartments dynamics. Data is normalized by the total number of 100kb regions considered in each sample. **D.** Scatter plot showing changes in compartment score between compartments defined at the control time point (Nut 0h) and after 1 hour of p53 activation (Nut 1h), Each dot represents a 100kb region. Dots are colored in black if the region contains at least one p53 binding site. Box plots represent the distribution of the compartment scores for each defined compartment dynamic, with or without the presence of p53. **E.** As in panel D, but in this case between compartments defined at the control time point (Nut 0h) and after 10 hours of p53 activation (Nut 10h). **F.** Distribution of differential compartment scores between compartments defined after 1 hour of p53 activation and control compartments (Nut 1h – Nut 0h). 100kb regions were classified according to their compartment dynamism and according to the presence or absence of functional p53 binding sites. Two-sample Wilcoxon test was performed to test whether differential compartment scores differed significantly between compartments which the same dynamics with and without functional p53 binding sites. **G.** As in panel F, but in this case differential compartments scores are between compartments defined after 10 hours of p53 activation and control compartments (Nut 10h – Nut 0h). **H.** Differential H3K27ac between 1 hour (Nut 1h) and 0 hour (Nut 0h) time points (log2FC), identified at 100kb regions (compartments) either with (p53) or without (control) an overlapping p53 binding site. Two-sample Wilcoxon test was performed to test whether p53 and control distributions differed significantly. **I.** As in panel H, but in this case analyzing the differential H3K27ac between 10 hours and 0 hours of p53 activation.

Collectively, these results show that direct p53 binding drives a displacement towards A compartments that supports the role of p53 acting solely as a transcriptional activator and suggest that indirect p53 effects, frequently occurring at later time points, are more heterogenous depending on the activating or repressing nature of downstream effectors (*e.g.,* p21^29^, E2F7^30^, and miRNAs^31^).

### Promoter interactome is highly rewired by p53 activation

Since TADs and DNA loops are thought to share a cohesin-driven loop extrusion formation mechanism^6,11^, we investigated whether our observed TAD dynamics during p53 activation may also be accompanied by promoter loop rewiring. To do so, we generate high quality and reproducible Promoter Capture Hi-C (PCHi-C) libraries using cells collected after 1 and 10 hours of Nutlin-3a treatment and control cells treated with the drug vehicle (0 hours) (Fig. 1A). Each sample was deep-sequenced, paired-end reads were mapped and filtered using the HiCUP pipeline, and significant interactions of 31,253 annotated promoters were called using the CHiCAGO pipeline (CHiCAGO score ≥ 5) (Fig. 4A and Supplementary Data 6). Specifically, we detected an average of 78,832 significant interactions per sample (SD = 4067), which were characterized by a median linear distance between promoters and their interacting regions of 260kb (SD = 12kb) and an average of 83% (SD = 2.5%) of promoter-to-non-promoter interactions (Supplementary Fig. 3A-B). PCA of CHiCAGO interaction scores demonstrated that promoter interactomes were highly reproducible within each condition and dynamic according to p53 activation state, which was also confirmed by hierarchical clustering (Fig. 4B and Supplementary Fig. 4C). Consistent with our previous Hi-C data, 1 hour of p53 activation led to the most dramatic changes in the promoter interactome. Specifically, early p53 activation triggered a loss of promoter interactions, mostly affecting longer interactions (> 200Kb) (Fig. 4C-E and Supplementary Fig. 4A), and an increase of promoter-promoter connectivity (Supplementary Fig. 4B). Late p53 activation (10 hours of Nutlin-3a treatment) also led to specific promoter-centric topologies, which included an increase of longer interactions (> 300Kb). A closer analysis allowed us to identify clusters of promoter interactions specifically gained at 1 hour (C1), 10 hours (C2) or both extensions of treatment (C7) of Nutlin-3a (Fig. 4F). Besides, we also recognized clusters of promoter interactions specifically lost during early (C3-4) and late (C6) p53 activation, some of these later re-established at 10 hours of Nutlin-3a treatment (C4).

**Fig. 4.**
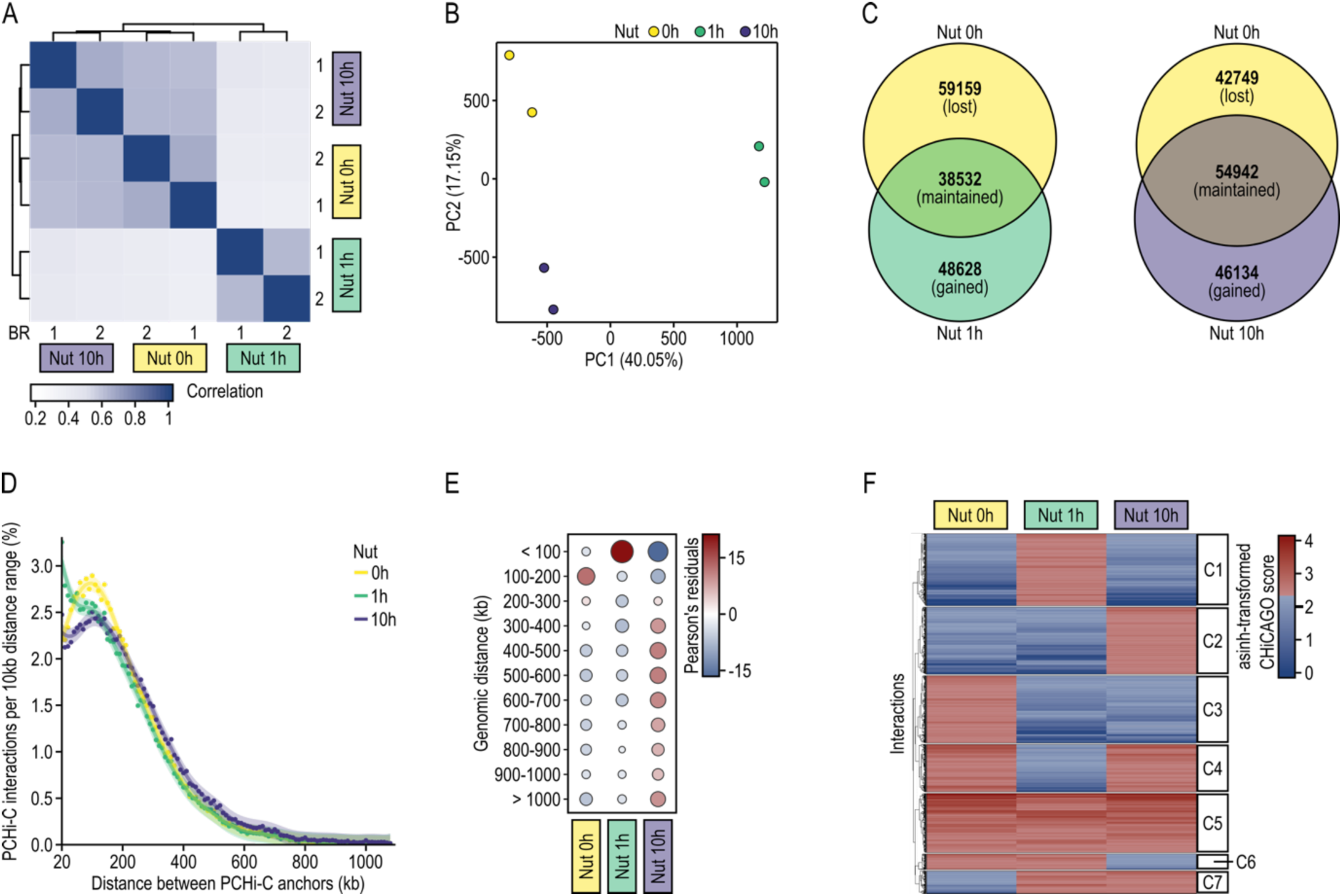
Promoter interactions shift during p53 activation. **A.** Pearson Correlation between promoter interactomes obtained by PCHi-C of biological replicates at Nut 0h, Nut 1h and Nut 10h time points. Samples were clustered hierarchically inputting Euclidean distances to the “complete” agglomeration method. Nut 0h refers to control cells without p53 activation. **B.** Principal Component Analysis of promoter interactomes throughout p53 activation obtained by PCHi-C at the biological replicate level. Numbers in parentheses represent the percentage of variance explained by each principal component (PC). **C.** Overlap between significant promoter interactions (CHiCAGO score ≥ 5) before and after 1 hour of p53 activation (left) and before and after 10 hours of p53 activation (right). **D.** Observed number of interactions with CHICAGO score ≥ 5 ranked according to the genomic distance between their anchor points. Distances were grouped into 10kb bins and converted to percentages to facilitate the comparison between samples. Observed percentages were fitted using a linear GAM (see methods). **E.** Correlation plot for resultant Pearson’s residuals from a Chi-square test of independence for the contingency table time-course *vs.* genomic distance bracket (X-squared = 2096.3, df = 20, p-value < 2.2e-16). Only significant associations are shown, where significance is determined as |Pearson’s residual| > 2. Bubble size indicates the number of interactions associated. Blue to red corresponds to an increasing association with genomic distance bracket. **F.** Heatmap of asinh-transformed CHiCAGO scores. Only interactions found significant in at least one time point were considered. Clusters were defined based on presence or absence of interaction significance in each time points, and hierarchical clustering was performed withing each predefined cluster to order the interactions order the clusters from C1-C7. Euclidean distances were measured between interactions and clusters, and the “complete” agglomeration method was used.

Summarizing, our data demonstrated that p53 activation leads to a dramatic rewiring of the promoter interactome that may contribute to reshaping the promoter-enhancer interactome landscape to ultimately control the transcriptional response.

### p53 controls transcription of previously non-associated genes located hundreds of kilobases away

Considering our results and previous knowledge, p53 acts preferentially as a distal regulatory element. In consequence, its target genes are often not directly identifiable, which accounts for a critical gap of knowledge. p53 regulation can occur from hundreds of kilobases away, bypassing several proximal genes, and can even occur in consequence to p53 binding in the intron of a non-target gene^32^. However, transcriptional regulation is conveyed through the physical proximity of communicating enhancers and promoters^33^. Therefore, we used our PCHi-C data to associate 253 p53-bound enhancers with 340 candidate genes to be distally (*i.e.,* over distance) controlled by p53, from now on referred to as p53 distal target genes (Fig. 5A-B and Supplementary Data 7). Among these, we identified examples of previously identified p53 target genes (*e.g., PPM1D*^34^*, TP53INP1*^35^*, PLK2*^36^*, DKK1*^37^*, FUCA1*^38^), but also identified potential new p53 target genes^39^ (*e.g., TGFR2*, *JAG2, BRD7, TENT4B, CD9*). p53 distal target genes were located at a median genomic distance of 116 kb from the functional p53 binding sites (Fig. 5C). Remarkably, only 7.1% (24/340) of these p53-bound enhancers were linked to the nearest gene, highlighting the importance of our long-range promoter interaction data to avoid misleading associations based on proximity within the genomic sequence. Indeed, unlike genes closest to the p53-bound enhancers along the linear genome (nearest genes, Fig. 5B), p53 distal target genes were enriched in p53-related gene sets (e.g., Reactome transcriptional regulation by TP53 pathway^40^, p53 dn.v1 up gene set^41^) (Fig. 5D and Supplementary Data 8).

**Fig. 5.**
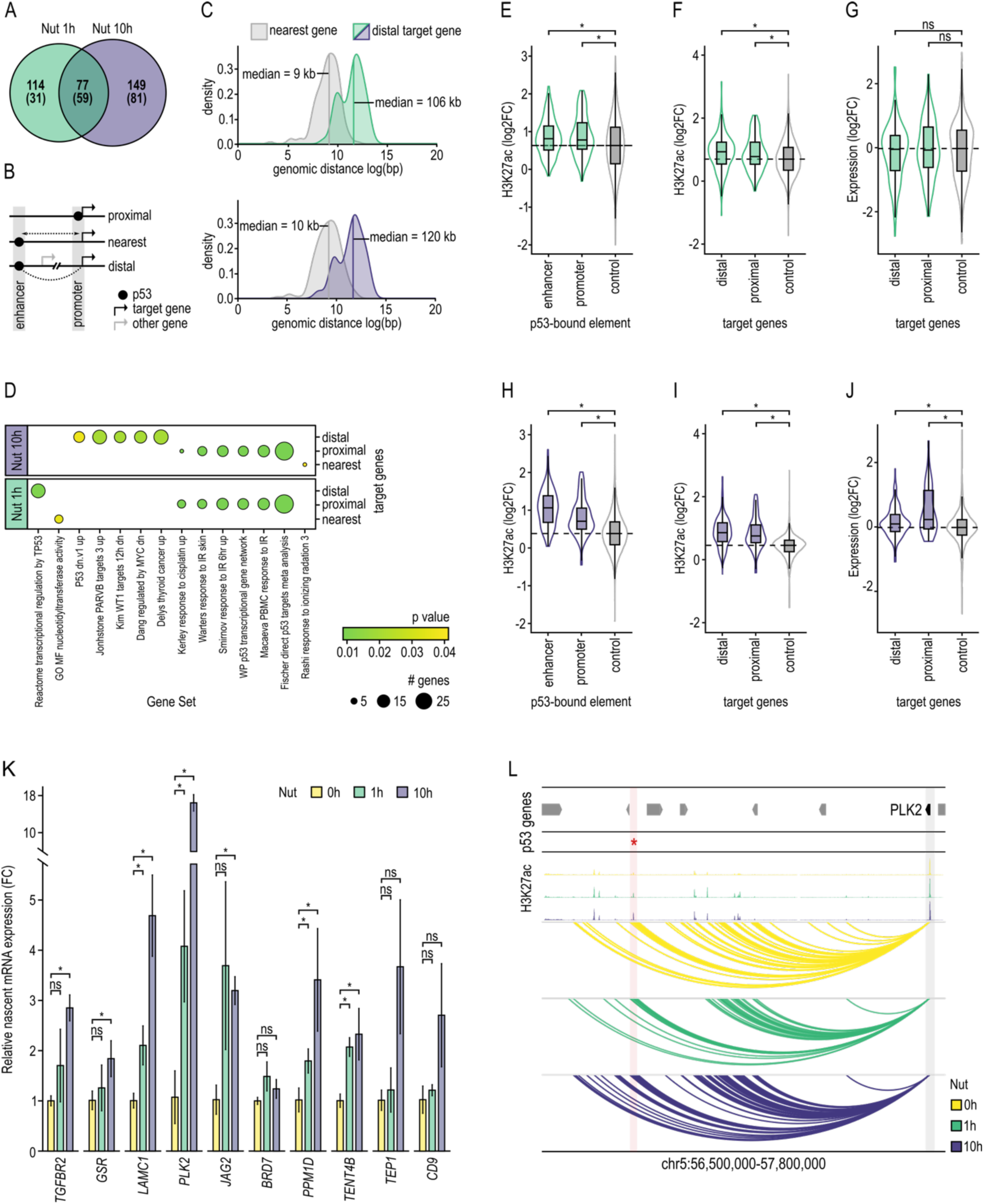
Identification of new p53 distal target genes through enhancer-promoter interactions. **A.** Overlap between p53 distal target genes at Nut 1h and Nut 10h time points. In parentheses are the number of p53 distal target genes which were primed (already interacting with a distal p53-bound enhancer prior to p53 activation) (*i.e.,* Nut 0h). Distal target genes are those genes engaging in PCHi-C interaction with a p53-bound enhancer. Nut 0h refers to control cells without p53 activation. **B.** Schematic representation of our definition of p53 target genes. Proximal target genes are those with p53 binding at their promoter. Nearest target genes are those found closest in linear proximity to a p53-bound enhancer. Distal target genes are those found to interact with a distal p53-bound enhancer (defined using PCHi-C data). **C.** Density distribution of the distance between the midpoint of a p53-bound enhancer and the transcription start site of the nearest gene (grey), or the transcription start site of the distal target gene linked by PCHi-C after 1 hour (green) or 10 hours (purple) of p53 activation. **D.** Gene set enrichment analysis of gene sets from the Molecular Signatures Database (MSigDB) with significant enrichment (p-adj < 0.05) in one or more p53 target gene groups. Bubble size indicates the number of genes found enriched for a given gene set, and green to yellow corresponds to the decreasing adjusted p-value for enrichment. p-values were calculated using a hypergeometric distribution test and adjusted for multiple comparison using Benjamini-Hochberg (BH). **E.** Differential H3K27ac between Nut 1h and Nut 0h time points (log2FC) at p53-bound regulatory elements (enhancer and promoter) and genome-wide (control). Genomic regions from p53-bound regulatory elements are excluded from genome-wide analysis. Mann-Whitney U-Statistic was used to test whether the mean H3K27ac log2FC differed significantly between groups using a p-value cutoff of 0.05 (star). **F.** Differential H3K27ac between Nut 1h and Nut 0h time points (log2FC) at promoter elements of p53 target genes, and promoter elements of non-targeted genes (control). Mann-Whitney U-Statistic was used to test whether the mean H3K27ac log2FC differed significantly between groups using a p-value cutoff of 0.05 (star). **G.** Differential expression between Nut 1h and Nut 0h time points (log2FC) of p53 target genes and non-targeted genes (control). Mann-Whitney U-Statistic was used to test whether the mean H3K27ac log2FC differed significantly between groups using a p-value cutoff of 0.05 (star). **H, I** and **J.** As in panels E, F and G, respectively, but considering target genes controlled by p53 at 10 hours of activation. **K.** Relative nascent mRNA expression (fold change) of p53 distal target genes along p53 activation. A two-tailed Student’s t-test was used to test whether relative expression differed significantly between adjacent time points for each gene (star). A linear mixed model was used to test whether relative expression differed significantly between adjacent time points, globally. For this, relative expression was averaged between the three replicates at each time point for each gene (p-value: 1e-8 after 1h, and 1e-7 after 10h. linear mixed model see methods). **L.** Interaction landscape of *PLK2’s* gene promoter (blue shade) showing stable interactions (arcs) with a p53-bound enhancer (red asterisk and pink shade) located 966kb downstream. H3K27ac profiles along p53 activation are represented at the top. Peak tips colored in black represent a peak going over the scale limit. Arrows symbolize gene placement and orientation along the genomic window.

Next, we validated these p53 distal target genes. First, we observed that enhancers and promoters bound by functional p53 binding sites gained significantly more H3K27ac than their control regions during p53 activation (Fig. 5E, H). Similarly, promoters of p53 distal target genes gained significantly more H3K27ac than genes that were not directly targeted by p53 (Fig. 5F, I). Moreover, these p53 distal target genes were upregulated at both the nascent transcribed (p-value: 1e-8 after 1h, and 1e-7 after 10h, linear mixed model see methods) and mature messenger RNA levels (Fig. 5G, J-K). Together, these results provide evidence of p53’s ability to drive transcription from a linear distal location. The *PLK2* kinase^36^ gene is an example of a p53 distal target gene, which is controlled by a p53-bound enhancer located 966kb downstream from its transcriptional start site (Fig. 5L). *PLK2’s* promoter and the p53-bound enhancer, both in spatial proximity facilitated by DNA looping, gained activity according to H3K27ac levels and nascent transcript (Fig. 5K-L). However, in this same example, genes closer to the p53-bound enhancer did not respond to p53 activation.

Altogether, our analysis demonstrates that DNA looping enables the effect of p53 activation to be broadcasted from enhancers to gene promoters and allows the identification of new p53 distal target genes to obtain a more complete picture of the p53 gene regulatory mechanisms.

### p53 binding at enhancers leads neo-loop formation with distal gene promoters

As we previously demonstrated, the promoter interactome was highly rewired during p53 activation (Fig. 4F). Next, we studied whether this rewiring could be caused, at least partially, by dynamic p53-mediated enhancer-promoter interactions. Only 47.1% (90/191) of p53 distal target genes at 1 hour of activation were primed (*i.e.,* gene promoter interacting with p53-bound enhancers prior activation) (Fig. 5A). This result suggests a novel role of p53 related to the establishment of DNA looping between enhancers and target gene promoters. Besides, just 23.6% (77/340) of distal target genes were in physical proximity with p53-bound enhancers at both time points (1h and 10h of Nutlin-3a treatment), most of which were primed (76.6% (59/77)).

Collectively, these results suggest the co-existence of two mechanisms by which p53 controls transcription of distal genes: i) a stable and primed mechanism that relies on a non-dynamic 3D chromatin structure, which could be associated with a core and stable transcriptional program; and ii) a dynamic and non-primed mechanism associated with the formation and destruction of DNA loops along p53 activation, which could enable tailoring of the response to different types of stress or DNA damage.

We then focused the study of the p53-mediated promoter-enhancer interactome rewiring at early p53 activation (*i.e.,* 1 hour of Nutlin-3a treatment) where p53-indirect contributions (*i.e.,* those related with factors transcriptionally controlled by p53) are minimal. Contrarily to the general trend of loss of interactions (Supplementary Fig. 4A), p53-bound enhancers increased connectivity with promoters located a median distance of 100kb away (Fig. 6A and Supplementary Fig. 5A-B). Specifically, 41.2% (91/221) p53-bound enhancers acquired interactions engaging distal target genes after 1 hour of Nutlin-3a treatment, none of these being engaged with other genes before p53 activation (Fig. 6B). On the other hand, 25.8% (57/221) of p53-bound enhancers lost connectivity with any gene promoters.

**Fig. 6.**
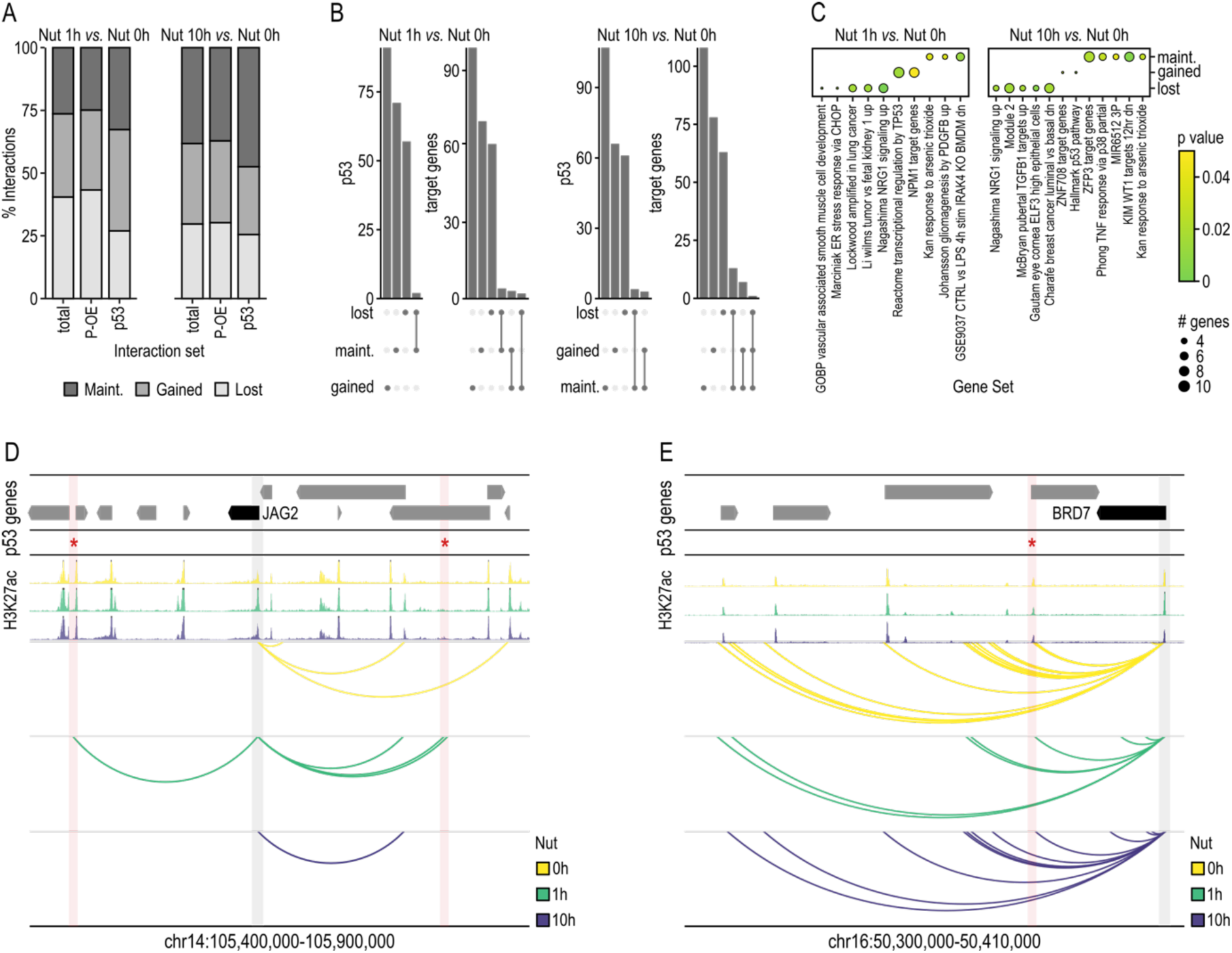
p53 drives the formation of promoter-enhancer interactions or uses pre- established ones to control gene transcription over distance. **A.** Percentage of significant promoter interactions (CHiCAGO score ≥ 5) gained, lost, and maintained between Nut 1h and Nut 0h time points (left) and Nut 10h, and Nut 0h time points (right) respectively. The three interaction sets shown are comparing the total interactomes (total); comparing promoter interactions with non-promoter regions (promoter with other-end or P-OE); and comparing interactomes with a p53 binding site present at either end (p53). Nut 0h refers to control cells prior to p53 activation. **B.** Upset plot showing the overlap between p53 binding sites and p53 distal target genes found at maintained, gained and lost interactions between Nut 1h and Nut 0h time points (left) and Nut 10h and Nut 0h time points (right). **C.** Gene set enrichment analysis of gene sets from the Molecular Signatures Database (MSigDB) with significant enrichment (p-adj < 0.05) in one or more p53 distal target gene groups defined by the interaction dynamism between Nut 1h and Nut 0h (left) and Nut 10h and Nut 0h time points (right). Bubble size indicates the number of genes found enriched for a given gene set. p-values were calculated using a hypergeometric distribution test and adjusted for multiple comparison using Benjamini-Hochberg (BH). **D.** Interaction landscape of *JAG2’s* gene promoter (blue shade) showing dynamic interactions (arcs) with two p53-bound enhancers (red asterisk and pink shade) located 186kb and 185kb away. H3K27ac profiles along p53 activation are represented at the top. Peak tips colored in black represent a peak going over the scale limit. Arrows symbolize gene placement and orientation along the genomic window. **E.** Interaction landscape of *BRD7’s* gene promoter (blue shade) showing stable interactions (arcs) with a p53-bound enhancer (red asterisk and pink shade). H3K27ac profiles along p53 activation are represented at the top. Peak tips colored in black represent a peak going over the scale limit. Arrows symbolize gene placement and orientation along the genomic window

Pathway-wise, genes that acquired *de novo* interactions with p53-bound enhancers minutes after p53 activation were enriched in the Reactome transcriptional regulation by TP53 pathway^40^ (*e.g., ATRIP*, *CARM1*, *DDIT4*, *GTF2H4*, *GSR*, *PRDX1*, *TCEB3*, *TP53INP1*, *YWHAH*) (Fig. 6C and Supplementary Data 7 and 8). Besides, we also identified other target genes not previously associated with p53 that are distally controlled by p53 through new loop (neo-loop) formation not observed in control cells previous to p53 activation (*e.g., E2F2*, *DUSP4*, *SMAD1*, *DNAJA3*, *PPM1D*, *YTHDC2*, *CHD1L*, *KMT2E*, *KIAA1429*). For example, *JAG2* gene, regulator of the Notch signaling pathway, is rapidly upregulated (*i.e.,* its promoter gains H3K27ac and its transcription is increased) by p53 via *de novo* formation of two DNA loops that engage one p53-bound enhancer each, located 186kb and 185kb away from *JAG2*’s transcriptional start site at 1 hour of treatment (Fig. 5K and 6D). This transactivation is transient since, at 10 hours of p53 activation, both loops are erased, its promoter lost H3K27ac and *JAG2* reduced its transcription rate.

This highly dynamic spatio-temporal promoter interactome contrasts with a topologically stable regulation of both well-known and novel genes by p53 (*e.g., CDKN2A, PLK2, GTF2F2*). For instance, the primed spatial proximity through stable DNA looping of the *BRD7* promoter and the p53-bound enhancer correlated with a gain in activity according to H3K27ac and an increase in nascent transcription levels during p53 activation (Supplementary Data 7 and Fig. 5K and 6E).

Collectively, these findings uncover an unexpected dependence of p53 on both newly formed and pre-existing enhancer-promoter loops to distally control gene transcription and ultimately determine the cellular response to cellular stress.

### Cohesin is required for the p53-mediated transcriptional response

As previously demonstrated, DNA looping enables long distance broadcasting of p53 transactivation from enhancers to gene promoters. On the other hand, previous studies have shown that cohesin plays a pivotal role in organizing spatio-temporal chromatin architecture by stabilizing DNA loops^42^. Given these premises, we next explored the dependency between p53 on cohesin to trigger a transcriptional response. To do so, we took advantage of HCT116 cells in which all alleles of the cohesin subunit *RAD21* were tagged with an auxin-inducible degron version 2 (AID)^43^. After validating RAD21’s rapid degradation after 6 hours of treatment with 1µM of 5-Ph-IAA and erase of DNA loops connecting p53-bound enhancers and distal target genes (Supplementary Data 6), we activated p53 using 10µM of Nutlin-3a (Supplementary Fig. 6A-B). Treated cells were used to: i) generate high quality ChIP-seq libraries to genome-wide profile the H3K27ac distribution, which allowed us to characterize active promoters (Fig. 7A-B and Supplementary Data 2); and ii) analyze nascent transcript by Real-Time Quantitative Reverse Transcription Polymerase chain reaction (qRT-PCR), which allowed the measurement of gene transcriptional activity.

**Fig. 7.**
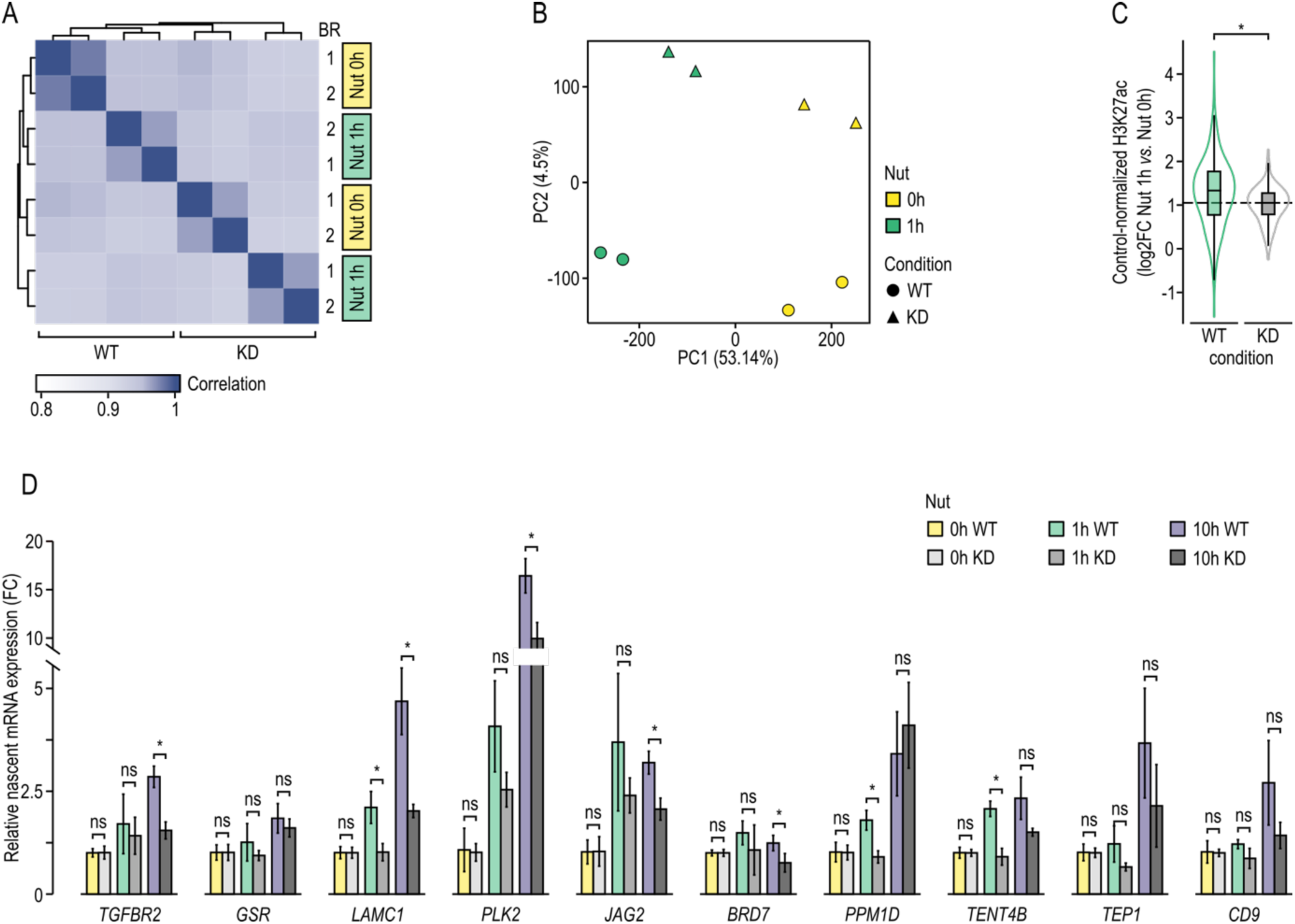
Cohesin depletion impedes p53-mediates transcriptional response. **A.** Pearson correlation between biological replicates (1 and 2) of H3K27ac ChIP-seq samples at time points Nut 0h and Nut 1h, both in wild type (WT) and knockdown condition of RAD21 (KD) using an auxin inducible degron system. Samples were clustered hierarchically using Euclidean distances and the “complete” agglomeration method. Nut 0h refers to control cells prior to p53 activation. **B.** Principal Component Analysis of H3K27ac ChIP-seq biological replicates of time point Nut 0h and Nut 1h both in wild type (WT) and knockdown condition of RAD21 (KD). Numbers in parentheses represent the percentage of variance explained by each principal component (PC). **C.** Control-normalized differential H3K27ac (log2FC) at promoters of p53 distal target genes between Nut 1h and Nut 0h time points in wild type (WT) and RAD21 depleted (KD) conditions. Control genes are defined as those not directly targeted by p53. Mann-Whitney U-Statistic was used to test whether the control-normalized mean H3K27ac log2FC differed significantly between groups using a p-value cutoff of 0.05 (star, p-value = 3.2e-6). **D.** Relative nascent mRNA expression (fold change) of p53 distal target genes along p53 activation in the presence (WT) or absence of RAD21 (KD). A two-tailed Student’s t-test was used to test whether relative expression differed significantly between conditions at each time point for each gene (star). A linear mixed model was used to test whether relative expression differed significantly between conditions of adjacent time points, globally. For this, relative expression was averaged between the three replicates at each condition and time point for each gene (p-value: 0.9 at Nut 0h, and 5e-9 at Nut 1h and 7e-6 at Nut 10h. linear mixed model see methods). **D.** Graphical representation of both mechanisms by which p53 controls transcription of distal genes: i) a stable and primed mechanism that relies on a non-dynamic 3D chromatin structure; and ii) a dynamic and non-primed mechanism associated with the formation and destruction of DNA loops along p53 activation.

Focused on the previously identified 340 p53 distal target genes (Fig. 5A-B and Supplementary Data 7), we observed that RAD21 degradation hindered H3K27ac gain at their promoters during early p53 activation compared with control genes, as opposed to the significant gain of H3K27ac observed in wild type conditions (Fig. 7C). In addition, we observed a significant drop in transcription rates depicted by the relative nascent mRNA expression levels (p-value: 0.9 at Nut 0h, 5e-9 at Nut 1h and 7e-6 at Nut 10h) (Fig. 7D and Supplementary Fig. 6C). For instance, RAD21 degradation impeded p53-mediated transcriptional upregulation of tumor suppressor gene *PLK2* ^36^ (Fig. 5K and 7D). Similarly, transcriptional upregulation of *LAMC1*, which encodes for a Laminin Subunit involved on cell adhesion, differentiation and metastasis^44^, as well as *TENT4B*, which encodes for a nucleotidyltransferase involved on RNA metabolic processes^45^, were also compromised (Fig. 7D).

Summarizing, these results disclose a novel association between p53 and cohesin in the context of cellular stress and demonstrate that RAD21 degradation reduces p53-mediated distal promoter activation and compromises p53-driven transcription response (Fig. 7E).

## Discussion

Although the tumor suppressor p53 was first characterized as a transcription factor 44 years ago ^46–56^, there are still many unresolved questions about its mechanism of action, including its interplay with the 3D genome organization to ultimately trigger the activation of hundreds of genes in a highly coordinated fashion. Here, using a fine-scale multi-omics time course, we demonstrated that p53 activation drives dramatic genome-wide changes in genome compartments, TADs and DNA loops to ultimately trigger a temporally dependent transcriptional response against cellular stress.

Unexpectedly, we observed that the most dramatic changes in spatio-temporal genome organization occur at 1 hour of p53 activation, prior to any dramatic p53 protein accumulation, significant cell cycle alterations or apoptosis. This finding agrees with a previous study that identified ∼200 genes being rapidly activated only 30 minutes after p53 activation using Global Run-on Sequencing (GRO-Seq) analysis^57^. However, with few exceptions^58–60^, most genome-wide alterations in 3D genome architecture associated with any cellular perturbation have been reported at later time points, suggesting a slower chromatin dynamism or advising the redesign of these time-resolved studies to capture early events.

These changes in spatio-temporal genome organization include direct p53 effects, immediately associated with its binding to DNA, and indirect p53 effects driven by factors transcriptionally regulated by p53. For instance, we have identified chromatin remodelers (*e.g., BRD7*, *CHD1L*), transcriptional co-activators (*e.g., CARM1*), transcription factors (*e.g., E2F2*, *SMAD1*), components and regulators of the transcriptional machinery (*e.g., GTF2H4*, *TCEB3, GTF2F2*) and epigenetic modifiers (*e.g., KMT2E*) as p53 distal target genes that may also contribute to DNA topological alterations. Direct p53 effects, which mainly contribute to dynamism at 1 hour of p53 activation, are primarily associated with gain of compartmental activity and enhancer-promoter loops. Here, we emphasize that these effects occur in early response, when the confounding effects of indirect transcriptional and post-transcriptional regulation are minimal. Our findings support the role of p53 as solely a transcriptional activator and demonstrate a previously unappreciated architectural role as a regulator at distinct topological layers. Contrarily, indirect p53 effects could mainly contribute to later dynamism and are more heterogenous depending on the activating or repressing nature of the downstream effectors.

The identification of reliable and reproducible p53 direct targets genes has been historically challenging due to the misleading association of p53-bound enhancers to target genes based on sequence proximity. Indeed, previous studies only considered as potential p53 direct targets those genes found differentially expressed following p53 activation or inactivation, whose transcriptional start sites are located from 5^61^ to 100 kb^62^ away from a p53 binding event. However, enhancers have the capacity to regulate genes located up to a few megabases away^63–65^ and p53-bound enhancers should not be an exception to the rule. To overcome this limitation, in this work we made use of PCHi-C^66^, a methodological breakthrough that we developed to associate distal regulatory elements and target genes in a genome-wide, unbiased manner^67,68^. Specifically, we identified 340 p53 distal target genes, including previously defined target genes as well as new ones. In fact, using as reference a survey of 346 target genes derived from 319 individual gene studies^39^ and a set of 3,509 target genes derived from 16 high-throughput data sets^39^, only 13 and 89 of our p53 distal target genes, respectively, were previously identified, in part due to the bias of analyzing only promoters and promoter-proximal regions. Indeed, p53 distal target genes that we identified were located at a median distance of 116 kb from the functional p53 binding sites, outside of the largest arbitrary window of 100kb previously established. Moreover only 7% of the p53 distal target genes that we identified were found to be the nearest gene of the targeting p53-bound enhancers. These results, all together, showcase the power of our long-range promoter interaction data to provide a more comprehensive landscape of the p53 direct transcriptional regulome. Another limitation in the identification of the true set of p53 target genes is the frequent use of late time points of p53 activation. This strategy unavoidably includes confounding indirect effects and does not capture early p53 targets genes dynamically regulated in time.

In this study we have also shown that the p53 regulome is characterized by extensive rewiring of the p53-bound enhancer-promoter loops. Previous studies^20–22,69^ proposed a mechanism, also called the permissive model, by which p53 relies on pre-programmed and stable enhancer– promoter loops to rapidly respond to p53 activating stimuli^70^. However, these studies used chromosome conformation capture technology to analyze only a few p53-bound enhancer-promoter loops. Using an unbiased genome-wide high-resolution approach, we have shown the co-existence of an additional dynamic and non-primed mechanism, called the instructive model, by which p53 controls transcription of distal genes through the formation and destruction of p53-bound enhancer-promoter loops (Fig. 7E). The permissive model could be associated with a rapid core transcriptional response structurally stable in time. Contrarily, the instructive model could be associated with a dynamic transcriptional response along time and may enable tailoring the response to different types of stress.

p53 binds a core set of regions regardless of cellular and epigenetic contexts via high affinity recognition of its binding motif without need of auxiliary transcription factors^26^. However, despite its largely invariant binding, p53 leads a highly cell-type-specific and stimulus-specific transcriptional response previously associated with permissiveness of the epigenetic environment^27,28^. On top of the dynamism along p53 activation, p53-bound enhancer-promoter loops could also be cell type and stimulus-specific, restricting the p53 targets available for direct transactivation. This type of structural guidance would ultimately allow tailoring the response to different types of stress responses and pathway activations. Further investigations to decode the cell-type-specific and stimulus-specific variation of the p53-centric spatio-temporal genome architecture will be needed to fully understand the variable p53 transcriptional response.

Emphasizing the relevance of spatio-temporal genome architecture, here we showed that cohesin was critically required for the induction of distal target gene expression triggered by p53 activation. Since cohesin removal leads to deletion of promoter-enhancer interactions^6,10^, our findings reinforce the critical role of DNA loops in mediating transcriptional changes in response to cellular stress. Paradoxically, cohesin depletion has little impact on steady-state gene transcription and enhancer activity^6^. However, two recent studies have implicated cohesin in inducible gene transcription. Specifically, it has been demonstrated that RAD21 removal hinders macrophages’ transcriptional response to the endotoxin lipopolysaccharide stimulation^18^ and prevents neurons from establishing transcriptional programs in response to new stimuli^19^. In agreement with these results, our findings demonstrate that cohesin is critical to support changes in transcriptional response, as opposed to the maintenance of a stable transcriptional landscape (Fig. 7E).

Although in this study we have been focused on the cohesin-dependent loop extrusion model, it is important to highlight that p53 has an intrinsically disordered domain^71^. Intrinsically disordered proteins have the capacity to undergo liquid-liquid phase separation to form membrane-less droplets that act as transcriptional condensates, promote gene expression, and remodel 3D chromatin organization^72^. However, to what extent p53 has the ability to induce topological changes in genome architecture through this mechanism and what the transcriptional consequences would be remains unknown.

Finally, our findings not only advance our understanding of p53 biology, but also pave the way for better-informed p53-based cancer therapies. Tumor suppressor gene *TP53* is the most frequently mutated gene in cancer^73^. p53-based therapy can either re-establish the functionality of mutant p53 proteins or safeguard wild type p53 from degradation in the cases of over-activity of its negative regulators (*e.g.,* MDM2, MDM4). However, only a few of these therapeutic strategies have reached late-stage clinical trials for various types of cancer^74–76^ and no drugs have been approved in Europe or USA up to date^77^. One of the multiple reasons behind the limited success of this strategy to safeguard wild type p53 could be the presence of loss-of-function mutations affecting p53 distal target genes. In this scenario, treatments to block p53 degradation would not be effective since the p53-mediated transcriptional response is compromised in a p53-independent manner. For this reason, a comprehensive list of true p53 direct target genes may help to better predict which patient will respond based on preservation of the wild type cascade downstream of p53. Another potential reason of failure of p53-based therapeutic approaches could be associated with mutations affecting cohesin. Although cohesin mutations are frequent in several types of cancer^73,78,79^, the mechanisms by which these mutations trigger cancer development and progression are not completely understood^80^. The Cancer Genome Atlas project has reported a comprehensive landscape of molecular alterations in bladder cancer, which includes mutations affecting p53 and cohesin in 49% and 11% of the cases, respectively^81^. Since our study disclose a p53 dependency on cohesin to drive a tumor suppressor transcriptional response, further investigations of mutually exclusive or synergistic effects of p53 and cohesin mutations are required to ultimately guide better therapeutic decisions.

Altogether, our study implicates spatio-temporal genome architecture as an instructive force for implementing a p53 transcriptional response to cellular stress, identifies a new set of reliable p53 direct target genes, and may help the future design of better-informed p53-based cancer therapies.

## Methods

### Cell culture

HCT116 or HCT116-RAD21-mAC cell lines were cultured in RPMI1640 medium with Glutamax (Fisher Scientific #61870044) supplemented with 10% FBS (Life Technologies #10270106) and penicillin-streptomycin (Dutscher #L0022-100) in a humid incubator at 37°C and 5% CO2. To activate p53 response, cells were supplemented with 10µM of Nutlin-3a (Quimigen #HY-10029) dissolved in DMSO and incubated for the required time. Degradation of the AID2-tagged RAD21 protein was induced by addition of 1µM of 5-Ph-IAA (MedChemExpress #HY-134653) dissolved in DMSO for at least 6h prior to further experimental procedure. Appropriate negative controls for each condition were established by incubating the cells with the drug vehicle (DMSO).

### Cell cycle analysis by FACs

After incubation with drugs or vehicle, 10^6^ cells per condition were collected in a single cell suspension, washed in PBS, fixed by adding 0.9ml of ice-cold 70% ethanol dropwise while vortexing and stored at -20°C for at least 24 hours. On the day of the analysis, cells were spun down at 1000g, 5 minutes and washed with PBS1x. Cells were stained for 30 minutes at room temperature using FxCycle PI/RNase Staining Solution (Life Technologies #F10797) following manufacturer’s instructions. Cell cycle was assessed using BD FACSCanto^TM^ II flow cytometer and data was analyzed using the Watson Pragmatic algorithm platform provided by FlowJo v10^82^.

### Western Blot

For validation of drug treatments, at least 5·10^6^ cells from different conditions were harvested, washed with cold PBS1x and lysed 30 minutes at 4°C in rotation, in ice-cold RIPA buffer (50mM TisHCl pH8, 150mM NaCl, 1%NP40, 0.5% Na-deoxycholate, 0.1%SDS) containing Complete, EDTA-free protease inhibitor cocktail (Merck Life Science #11873580001), 1mM PMSF (Merck Life Science #10837091001) and 1mM DTT (Merck Life Science #DO632- 1G). Cell lysates were sonicated using a UP50H Hielscher sonicator (1cy, 90%amp, 10 seconds per burst, 3 bursts per sample) and clear cell extracts were collected after centrifugation at max speed, 8 minutes, 13°C. Total cell lysates were quantified using BCA Pierce^TM^ (Thermo Scientific #23225) and at least 20µg of protein were resolved on 8 or 10% polyacrylamide gels, transferred onto nitrocellulose membranes and incubated with appropriate primary and secondary antibodies. The following primary antibodies were used: p53 DO1 (Santa Cruz Biotechnology, sc-126, diluted 1:200), RAD21 (Abcam, ab992, diuted 1:1000), Vinculin (Abcam, ab129002, diluted 1:10000), α-Tubulin (SIGMA-ALDIRCH, T6199, diluted 1:10000). The following secondary antibodies were used: IRDye 800CW Goat anti-rabbit IgG secondary antibody (LICOR #926-32211, dilute 1:10000) and IRDye 680RD Goat anti-mouse IgG secondary antibody (LICOR #926-68070, dilute 1:10000). Immunoblots were imaged using an Odyssey CLx imager and Image Studio Lite v5.2

### *In situ* Hi-C and PCHi-C libraries generation

Hi-C and PCHi-C libraries were prepared as described in^69^. Briefly, cells were fixed in DMEM medium supplemented with 10% FBS and 2% methanol-free formaldehyde for 10 minutes rotating at room temperature. After quenching the formaldehyde with 0.125M glycine and washing the cells with 1X PBS, nuclei were extracted by lysing the cells in 10 mM Tris-HCl pH 8.0, 10 mM NaCl, 0.2% IGEPAL CA630 (SIGMA-ALDRICH #18896-50ML), 1× cOmplete EDTA-free protease inhibitor cocktail. Chromatin was digested overnight with HindIII enzyme and the cohesive restriction fragment ends were filled-in with dCTT, dTTP, dGTP and biotin-14-dATP (Invitrogen #19524-016) nucleotides. After blunt-end ligating the restriction fragment ends, the chromatin was decrosslinked by incubating overnight with proteinase K and purified by phenol:chloroform:isoamyl alcohol 25:24:1 (SIGMA-ALDRICH #P3803-100ML) extraction.

Non-informative biotin at restriction fragment ends was removed by incubating the samples with dATP and T4 DNA polymerase (New England Biolabs #M0203S). After purifying the DNA again with a phenol:chloroform:isoamyl alcohol extraction, 10µg of the chromatin was sheared using a Covaris M220 focused ultrasonicator in 130µl cuvettes using the following parameters: 20% duty factor, 50 peak incident power, 200 cycles per burst, 65 seconds. The ends were end-repaired and dATP-tailed, followed by a biotin-pulldown using Dynabeads MyOne streptavidin C1 paramagnetic beads (Thermo Fisher #65001) to enrich for those DNA fragments which contain information of a chromatin loop. PE Illumina adapters (Supplementary Data 9) were ligated to the sample and the library was amplified for 8 cycles. Finally, the library was purified using a SPRI bead double-sided selection (0.4-1 volumes). Size and concentration of the finished Hi-C libraries was assessed by DNA ScreenTape Analysis (Agilent #5067-5582) on an Agilent 2200 Tapestation.

For PCHi-C libraries, 500-1000ng of Hi-C library were captured using the SureSelectXT Target Enrichment System for the Illumina Platform (Agilent Technologies) as instructed by the manufacturer. Captured library was amplified a total of 4 cycles, and the size and concentration of the finished PCHi-C libraries was assessed by high-sensitivity DNA ScreenTape Analysis (Agilent #5067-5584) on an Agilent 2200 Tapestation.

### ChIP-seq library preparation

Cells were crosslinked in 1X PBS supplemented with 1% methanol-free formaldehyde (Thermo Fisher #28908) for 10 minutes rotating at room temperature. After quenching the formaldehyde with 0.125M glycine for 5 minutes rotating at room temperature. After washing the cells with 1X PBS, pelleted cells were lysed in 1% SDS (AppliChem #A0676,0250), 10mM EDTA (Invitrogen #AM9260G), 50mM Tris-Cl pH 8’1 (Invitrogen #AM9855G) at 4°C for 20 minutes. Samples were sonicated using a Covaris M220 focused ultrasonicator at a concentration of 20·10^6^ cells/ml using the following parameters: 10% duty factor, 75 peak incident power, 200 cycles per burst, 15 minutes. After centrifuging the samples at 14000rpm to remove cell debris and recovering the chromatin-containing supernatant, 33µl of sonicated chromatin were prepared for immunoprecipitation by adding 267µl of buffer containing 1% Triton (AppliChem #A4975,0100), 1’2mM EDTA, 16’7mM Tris pH 8, 167mM NaCl (Invitrogen #AM9760G), 1X cOmplete protease inhibitor cocktail (Merck cat. #11873580001) and the appropriate amount of antibody (1ug of α-H3K27ac and 0.5ug of α-H3K4me1; Diagenode #C15410196 and #C15410194 respectively). Samples were incubated rotating at 4°C overnight. Chromatin was immunoprecipitated by adding 10µl of both protein A and protein G-conjugated paramagnetic beads (Invitrogen #1001D and #1003D, respectively) and incubated at 4°C for 1h. After washing the beads, chromatin was decrosslinked overnight at 65°C using proteinase K (ThermoFisher Scientific #EO0491) and purified using 1.1 volumes of SPRI beads according to the manufacturer’s instructions (CleanNA #CNGS-0050). For ChIP inputs, an equivalent amount of sonicated chromatin was directly decrosslinked and purified as before.

ChIP-seq libraries were performed using the KAPA HyperPrep Kit (Roche #07962363001) according to the manufacturer’s instructions using Truseq Illumina adapters and PCR primers described in Supplementary Data 9. Samples were sequenced to reach a minimum number of either 20M or 45M valid paired-read for H3K27ac and H3K4me1 histone marks respectively.

### RNA-seq library preparation

RNA was extracted from 200,000 frozen cell pellets using the RNAeasy Mini Kit (Qiagen #74104) following the manufacturer’s instructions. Total RNA’s integrity and concentration was assessed using RNA ScreenTape Analysis (Agilent #5067-5576) on an Agilent 2200 Tapestation. Samples were sequenced to reach a minimum number of 30M unique valid paired reads.

### qRT-PCR

RNA was extracted from the corresponding conditions from 200,000-250,000 cell pellets using the RNeasy Mini Kit (Qiagen #74104) according to the manufacturer’s instructions. On-column DNase (Qiagen #79254) treatment was applied to ensure no traces of genomic DNA were carried over during the genomic purification. The RNA was retrotranscribed using the SuperScript III First-Strand Synthesis System kit (ThermoFisher #18080051) according to the manufacturer’s instructions using random hexamer priming and the final cDNA was further diluted to 3.7 ng/ul.

qRT-PCR primers for a given gene were designed against the most common intron between alternative transcripts (merged Ensembl/Havana database for protein coding transcripts) using Primer3 web tool (v.4.1.0 https://primer3.ut.ee/) using the following parameters: product size ranges 100-150, optimal primer Tm 60°C, the rest in default. The primers were further checked for unique hybridization on the genome using the UCSC BLAT web tool using the GRCh37/hg19 human version of the genome, and tested for their amplification efficiency (between 85-115%, Supplementary Data 9).

The RT-qPCR reactions were carried out on 384-well plates using a QuantStudio 7 Flex Real-Time PCR System (ThermoFisher #4485701) using 10 µl reactions (2 µl of cDNA 3.7 ng/µl, 0.5 µl of forward and reverse primer mix 10 µM each, 5 µl of SYBR Green PCR Master Mix (ThermoFisher #4368577) and 2.5 µl of nuclease-free water). Raw data was analyzed using the QuantStudio software (v.1.3) and the nascent transcript expression fold change was calculated using the ΔΔCt method^83^ against *HPRT1* as the housekeeping gene. Primer sequences are de- tailed in Supplementary Data 9, and nascent transcript expression fold change data is summa- rized in Supplementary Data 10.

We adjusted a linear mixed model to relative nascent mRNA expression to decide whether gene expression can be explained by the fixed-effect of the experimental condition (either the effect of Nutlin-3a over time or degradation of RAD21), or by a random effect. p-values correspond to the rejection of a hypothesis which states that the random effect fits better that the experimental condition.

### Hi-C processing

Various steps were taken to process the Hi-C data, including read quality control, mapping, interaction detection, filtering, and matrix normalization using the TADbit pipeline^23^ (specific version: https://github.com/fransua/TADbit/tree/p53_javierre). The mapping of di-tags was performed using GEM3 mapper^84^ onto the GRCh37.p13 reference genome (hg19 downloaded from UCSC, http://genome.ucsc.edu). TADbit mapping consists in two steps. First full reads are mapped, then, for the remaining unmapped reads, TADbit searches for HindIII ligation sites consisting of facing fragments of restriction-enzyme (RE) sites (*e.g.,* AAGCTAGCTT). These reads with ligation sites are then split and their original HindIII site is reconstructed. Second, each of the read fragments undergo a second round of mapping. This methodology may result in reads mapped in more than the two locations expected for Hi-C di-tags. TADbit crumbles these multiple mappings (see Supplementary Data 1) into multiple individual di-tags.

Di-tag filtering was then conducted on pairs of mapped read fragments in order to remove experimental or computational artifacts. Filters used in TADbit were 1, 2, 3, 4, 6, 7, 9, and 10 (see TADbit online documentation); namely we filtered for di-tags in the same RE-fragment (1: self-circle, 2: dangling-end, and 3: mapping-errors); or in contiguous fragments (4: extra dangling-end); for di-tags mapped in uninformative RE fragments, either too short, below 50 nucleotides, or too large, above 100kb (6: too short, 7: too large); for PCR artifacts (9: duplicated); and finally, we also filtered for di-tags with at least an end mapped too far from any RE-site, as Hi-C product should necessary come from their vicinity (10: random breaks). The specific distance considered to fall in this last category is automatically defined by TADbit using the length distribution of single-fragment di-tags (specifically di-tags falling in the second filer “dangling-end”).

Hi-C interaction matrices at 50kb and 100kb resolutions were generated from filtered di-tags. These matrices are built, for each time point, by combining biological replicates. Interaction matrices were then cleaned by removing bins (rows or columns) with a ratio of mid-range cis interactions (interactions below 5Mb) over total interactions below 1. Bins with more than expected long-range interactions were considered artefactual. The genomic interaction matrix was normalized using the ICE normalization as defined in^85^.

A/B compartments were called independently for each chromosome and time point at a 100kb resolution. To this end, ICE-corrected interaction matrices were further distance corrected and finally transformed into Pearson correlation matrices^86^. In order to reduce potential noise in our data matrix we used a median filter with a size of 3 bins, similar to the methodology implemented in the HOMER software^87^. For each chromosome, we computed the three leading eigenvectors and performed a manual assessment to determine which one potentially provided a more accurate representation of the heterochromatin and euchromatin segregation. The parameters considered for the decision of the eigenvector to use were: the correlation with GC-content (expected to be high in A-compartments), the enrichment of our available markers for activity (H3K27ac and RNA-seq), the relative importance (eigenvalue) of each of the eigenvectors, the correlation between time points, and the general pattern and distribution of compartments in the interaction matrices. As expected, we selected the first eigenvector as representative of compartment segregation for most of the chromosomes in all time points. However, in the cases of chromosomes 4 and 7, we selected their second eigenvectors, this again, for all time points. Following the accepted protocol, the selected eigenvectors were rotated according to their enrichment in markers for transcriptional activity in order to associate positive values to A-compartment and negative values to B-compartments. Finally, a sigmoid transformation was applied to the resulting eigenvector in order to reduce the impact of specific outliers across time points and chromosomes. We referred to the resulting transformed eigenvectors as compartment scores.

TAD borders were identified using the TADbit built-in TAD caller applied on 50kb interaction matrices. TADbit’s TAD caller assesses the robustness of the TAD border detection by assigning a score between 1 and 10 to each TAD border. In this work, we used only TAD borders with a score strictly above 4. As an alternative TAD calling strategy, we also used TAD-borders called using the insulation score as proposed in^24^. In our implementation, we measured levels of interactions in the distance range of 50kb to 400kb and considered a value of *delta=2* (smoothing parameter, where 2 relates to the span in number of bins). This combination of parameters was found by maximizing the number of shared borders between replicates. TAD border alignments were generated with the function *align_TAD_borders* from our version of TADbit. We considered borders to be homologous between replicates if the distance between them was below or equal to 100kb (2 bins).

### ChIP-seq processing

ENCODE standards were followed to process paired-end reads. Sequencing adapters were trimmed using Trim Galore! (0.6.6). Reads were then mapped to the reference genome (GRCh37.p13) using bowtie2 (2.3.2)^88^ in the *--very-sensitive* mode. Low-quality reads, reads overlapping the ENCODE blacklist, and duplicate reads were filtered out using samtools (1.9)^89^. Genome-wide coverage was computed using the function bamCoverage from deepTools (3.2.1)^90^ to obtain bigwig files for visualization purposes. Macs2 (2.2.7.1)^91^ was used for peak calling in the narrow mode for H3K27ac and broad mode for H3K4me1, using an input sample as control, with default parameters. Consensus peaks were computed for each condition using Macs2 with all replicates and their respective input samples as control, setting the parameter *--scale-to small*. For quantification, a set of non-redundant enriched regions were defined by taking the union of peaks from all datasets. The signal of this set of regions was then quantified for each sample by counting the number of reads falling into each region using the function regionCounts from R package csaw (1.30.1)^92^. Normalization and differential analysis were performed using R package DESeq2 (1.36.0)^93^. Library size factors for normalization were calculated based on the background signal. This background signal was quantified per sample, excluding the set of previously defined non-redundant enriched regions. Specifically, this signal was quantified on fragment counts over genomic bins of 10kb using the function windowCounts from R package csaw^92^. Library statistics were assessed using FastQC and MultiQC^94^ and summarized in Supplementary Data 2.

### RNA-seq processing

ENCODE standards were followed to process paired-end reads. Sequencing adapters were trimmed using Trim Galore! (0.6.6). Reads were then mapped to the reference genome (GRCh37.p13) using STAR (2.7.0f)^95^ with parameters recommended by ENCODE. Read counts were quantified using featureCounts from subread (2.0.0)^96^. Normalization and differential analysis were performed using R package DESeq2 (1.36.0)^93^. Library statistics were assessed using FastQC and MultiQC^94^ and summarized in Supplementary Data 3.

### Definition of promoters and enhancers

Gene promoters were defined as regions spanning 1,000 base pairs upstream and 200 base pairs downstream from their transcriptional start site. Ensembl gene annotation GRCh37 release 87 was used to define transcriptional start sites. To define enhancers, we first identified the intersection between consensus H3K4me1 and H3K27ac peaks at 0 hours, 1 hour and 10 hours after p53 activation in wild type conditions. Consensus enhancers were defined as the union of enhancers at each time point. To define enhancer activity, we annotated whether consensus H3K4me1 and H3K27ac peaks were present for each time point, with solely H3K4me1 presence signifying a primed enhancer state, and both H3K4me1 and H3K27ac presence signifying an active enhancer state. Defined enhancer regions used for downstream analysis are presented in Supplementary Data 4.

### Definition of functional p53 binding sites

We used publicly available ChIP-seq data of p53 in the HCT116 cell line treated with Nutlin- 3a from^25^ (GSE86164). This dataset was refined into time point-specific functional p53 binding sites by intersecting it with our H3K27ac consensus peaks dataset. Specifically, the H3K27ac consensus peaks of 1 and 10 hour time points in wild type conditions were overlapped with the p53 binding sites using R package GenomicRanges (1.50.2)^97^. A p53 binding site was defined as functional when overlapping at least 1 base pair with an H3K27ac peak. The set of functional p53 binding sites are listed in Supplementary Data 5.

### PCHi-C processing

Raw sequencing reads were processed using HiCUP (0.8.2)^98^. The target sequence of the restriction enzyme HindIII was used to computationally digest the genome. HiCUP was then used to map paired-end reads to the human genome (GRCh37.p13), filter out experimental artifacts such as circularized reads and re-ligations, and remove duplicate reads. Paired reads which do not overlap with a captured restriction fragment are filtered out, retaining only uniquely captured valid reads for downstream analysis. This information is then used to assess the sample’s capture efficiency. Datasets were scaled down to the same sequencing depth in reference to the corresponding biological replicate of the dataset with the lowest number of unique valid reads (the 1 hour time point datasets). Interaction confidence scores were computed using the R package CHiCAGO (1.24.0)^99^. Interactions with a CHiCAGO score ≥ 5 were considered high-confidence interactions. CHiCAGO scores were recalibrated based on control datasets (0 hours) using the fitDistCurve function from the R package CHiCAGO. Library statistics were assessed using FastQC and MultiQC^94^ and are presented in Supplementary Data 6.

Linear generalized additive model (linear GAM) was used to fit percentages of observed significant interactions at different genomic distance brackets with splines. For all samples, the residuals calculated were all above 0.99 and the p-values associated with the number of parameters below 1e-16, justifying the optimized (grid-search) choice of 21, 31 and 30 splines for 0 hour, 1 hour and 10 hour time points, respectively (smoothing lambda parameter optimized together at 0.1 for all three samples). We used the GAM implementation from pyGAM^100^. Prediction intervals were calculated from the distribution of observed data points. Confidence intervals of the model, not shown, were fully included inside prediction intervals.

### Definition of p53 target genes

Proximal p53 target genes are defined as those genes with an overlapping functional p53 binding sites at their promoter. Distal p53 target genes are determined based on high-confidence interactions defined using CHiCAGO (1.24.0)^99^ (CHiCAGO score ≥ 5). These distal genes must be located in captured fragment interacting with a fragment that overlaps a functional p53 binding site that i) is not defined as proximally targeting, and ii) overlaps an active enhancer. For each p53 binding site, a prioritization strategy was implemented whereby the distal target gene with the largest gain in H3K27ac at their promoter (based on the mean log2 fold change) was prioritized as their target gene. Genes targeted by the same p53 binding site and with the same mean log2 fold change were all defined equally as distal target genes. The set of p53 target genes is summarized in Supplementary Data 7.

### Gene set enrichment analysis

Gene set enrichment analysis (GSEA) was performed using the enricher function from clusterProfiler (4.4.4)^101^ using default parameters. p-values were computed using a hypergeometric distribution test and adjusted for multiple comparisons using Benjamini-Hochberg correction. Gene sets were defined as enriched with an adjusted p-value ≤ 0.05. Gene sets were obtained from the Molecular signatures database^101^ via the msigdbr R package (7.5.1). Analysis was performed against all Human Collections, with particular emphasis on H collection (hallmark gene sets)^102^, C2 collection (curated gene sets), C5 collection (ontology gene sets)^103–105^ and C6 collection (oncogenic signature gene sets). Depending on the analysis, we tested given sets of genes against different libraries of total genes (often referred to as a gene universe). For the GSEA of RNA-seq and ChIP-seq differential analysis results, we used the Ensembl gene annotation GRCh37 release 87. For the GSEA of p53 target genes, we used genes present captured HindIII fragments. All GSEA results are presented in Supplementary Data 8.

### Statistical methods and figures

PCA of TAD borders and compartments was performed using the scikit-learn API v1.1.3. Statistical tests and complex numerical treatments were performed with SciPy v1.10.1^106^ and NumPy v.1.24.2^107^. Matrix comparison and large internal data manipulation were performed using the Pandas API. Figures and plots were generated using ggplot2 (Wickham H (2016). *ggplot2: Elegant Graphics for Data Analysis*. Springer-Verlag New York. ISBN 978-3-319- 24277-4) and matploltib v3.7.1^108^.

PCA for RNA-seq, ChIP-seq, and PCHi-C samples were generated using prcomp function from stats (4.2.1) R package^109^. Statistical tests were performed using ggpubr (0.6.0)^110^. Data manipulation and complex numerical treatments were performed using tidyverse (1.3.2).

Weighted Euclidean distances were applied on principal components (PCs) computed from Hi-C data to measure the similarity of biological replicates. The formula applied was:

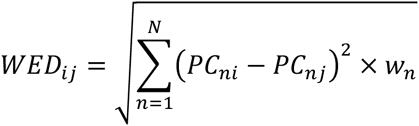

where *i* and *j* are biological replicates, *N* the number of considered principal components (10 here), and *w_n_* the variance explained by the current PC, and *PC_ni_* the current (*n*) PC of the sample *i*.

For analysis regarding the identification of p53 target genes, mean comparisons were performed using a Mann-Whitney U-Statistic. Data manipulation and integration with PCHi-C data sets was performed using HiCaptuRe^111^.

We implemented the HiCRep algorithm^112^ in TADbit to compute the stratum-adjusted correlation coefficient (SCC) between the two replicates.

### Data visualization

Data from this study can be visualized in the Washington University Epigenome Browser (http://epigenomegateway.wustl.edu/browser/)

## Data availability

Raw and processed data sequencing data for Hi-C, ChIP-seq, RNA-seq and PCHi-C have been deposited in Gene Expression Omnibus (GEO) under the accession number X.

## Code availability

Scripts to reproduce the analysis and figures in this study are available on GitHub https://github.com/JavierreLab/p53.

## Acknowledgements

We gratefully acknowledge Masato T. Kanemaki and Sergi Cuartero for kindly sharing his HCT116 RAD21-mAID-mCloverCMV-OsTIR1(F74G) cell line model^43^ and his antibody against RAD21 respectively. We thank Joaquin M. Espinosa team for critical discussion about publicly available RNA-seq datasets. Finally, we want also to thank Wouter de Laat and members of the Javierre Group for critical discussion. We thank CERCA Programme/Generalitat de Catalunya and the Josep Carreras Foundation for institutional support. This work was supported by FEDER/Spanish Ministry of Science and Innovation (PID2021-125277OB-I00), the European Hematology Association (4823998) and the Scientific Foundation of the Spanish Association Against Cancer (AECC) (LABAE21981JAVI). BMJ is funded by the Spanish Health Institute Carlos III (CP22/00127), LR is funded by an AGAUR FI fellowship (2019FI-B00017). The funding bodies were not involved in the study design, collection, analysis, interpretation of data, the writing of this article or the decision to submit it for publication.

## Author contributions statement

FS, LF and BMJ designed research and supervised the project; LF, LR, AL, BU, AA, EMN and BMJ performed research; FS, AN-A, MC analyzed data; ALO, PF, MG, SDC, JLS and AV provide critical feedback; BMJ wrote the paper and all authors provided feedback on the manuscript.

---

**Suppl. Fig. 1.**
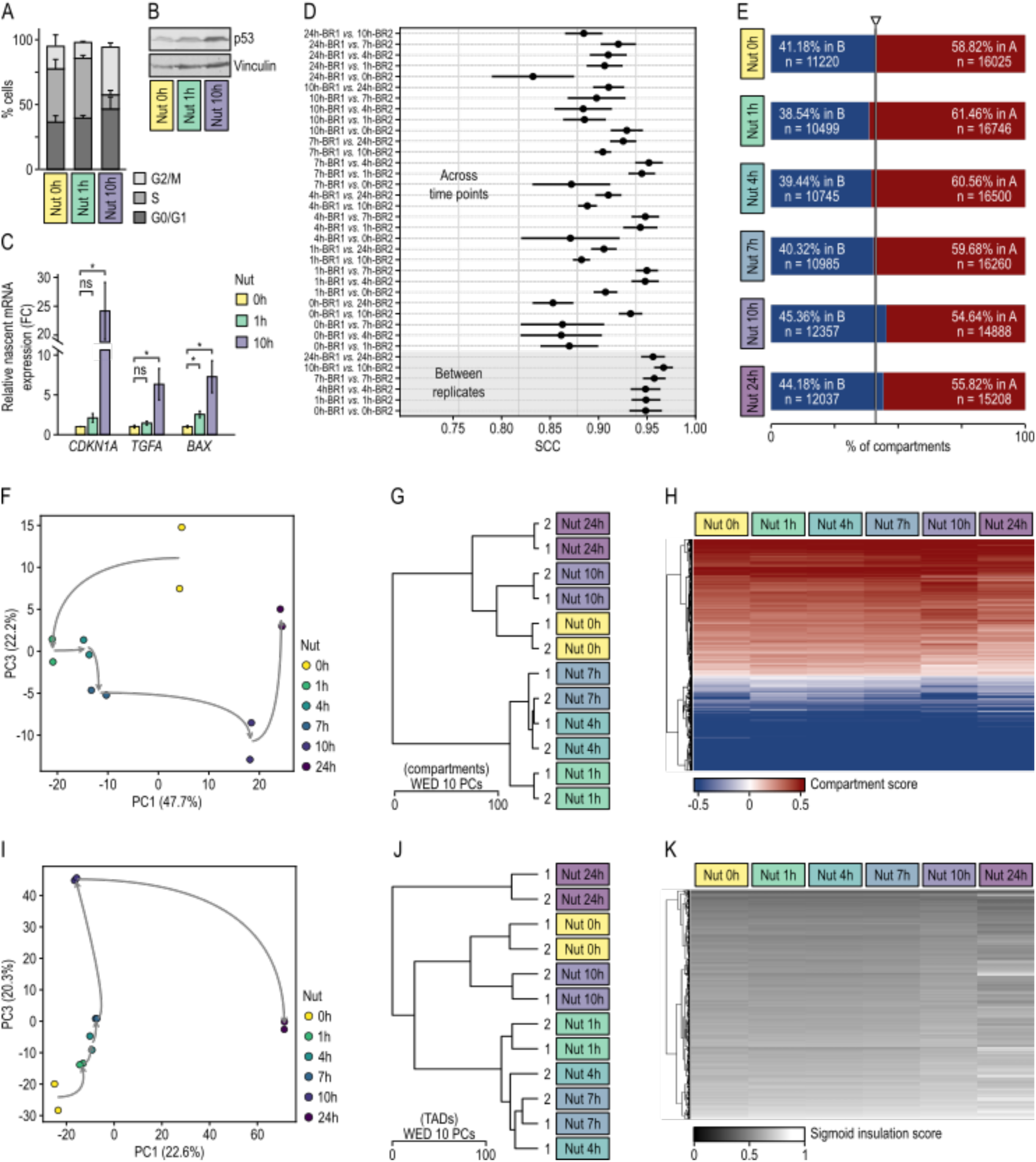
**A.** Representative distributions of cells in each cell cycle phase (G0–G1, S, G2–M) during Nutlin-3a treatment. 0h means control cells without p53 activation. **B.** Western blot of p53 at 0, 1 and 10 hour timepoints, using Vinculin as control. **C.** Levels of relative nascent mRNA expression along p53 activation for 3 known p53 target genes, measured by qRT-PCR. **D.** Stratum Adjusted Correlation Coefficient (SCC), with standard deviation bar, between time points and replicates of Hi-C data. **E.** Percentage of 100kb genomic regions defined as A or B compartments for each time point. **F.** Principal Component Analysis of the compartment scores for all Hi-C biological replicates throughout p53 activation. Principal components 1 and 3 were taken into consideration. Numbers in parentheses represent the percentage of variance explained by each principal component (PC). **G.** Hierarchical clustering analysis (Ward criterion) of compartment values throughout p53 activation reflecting the degree of dissimilarity between biological replicates of different samples. The distances between replicates were measured by applying weighted Euclidean distance (WED) of the 10 first principal components. **H.** Compartment scores of the non-dynamic compartments upon p53 activation. Each row represents a 100kb bin and each column represents a time point. **I.** Principal Component Analysis of TADbit TAD border scores for all Hi-C biological replicates throughout p53 activation. Principal components 1 and 3 were taken into consideration. Numbers in parentheses represent the percentage of variance explained by each principal component (PC). **J.** Hierarchical clustering analysis (Ward criterion) of TADbit TAD border scores across p53 activation reflecting the degree of dissimilarity between biological replicates of different samples. The distances between replicates were measured by applying weighted Euclidean distance (WED) of the 10 first principal components. **K.** Sigmoid-normalized TAD insulation scores of the TAD borders that remained unchanged across all samples (invariant TAD borders). Color intensity represents insulation score, with lower scores (in black) indicating stronger insulation between TADs.

**Suppl. Fig. 2.**
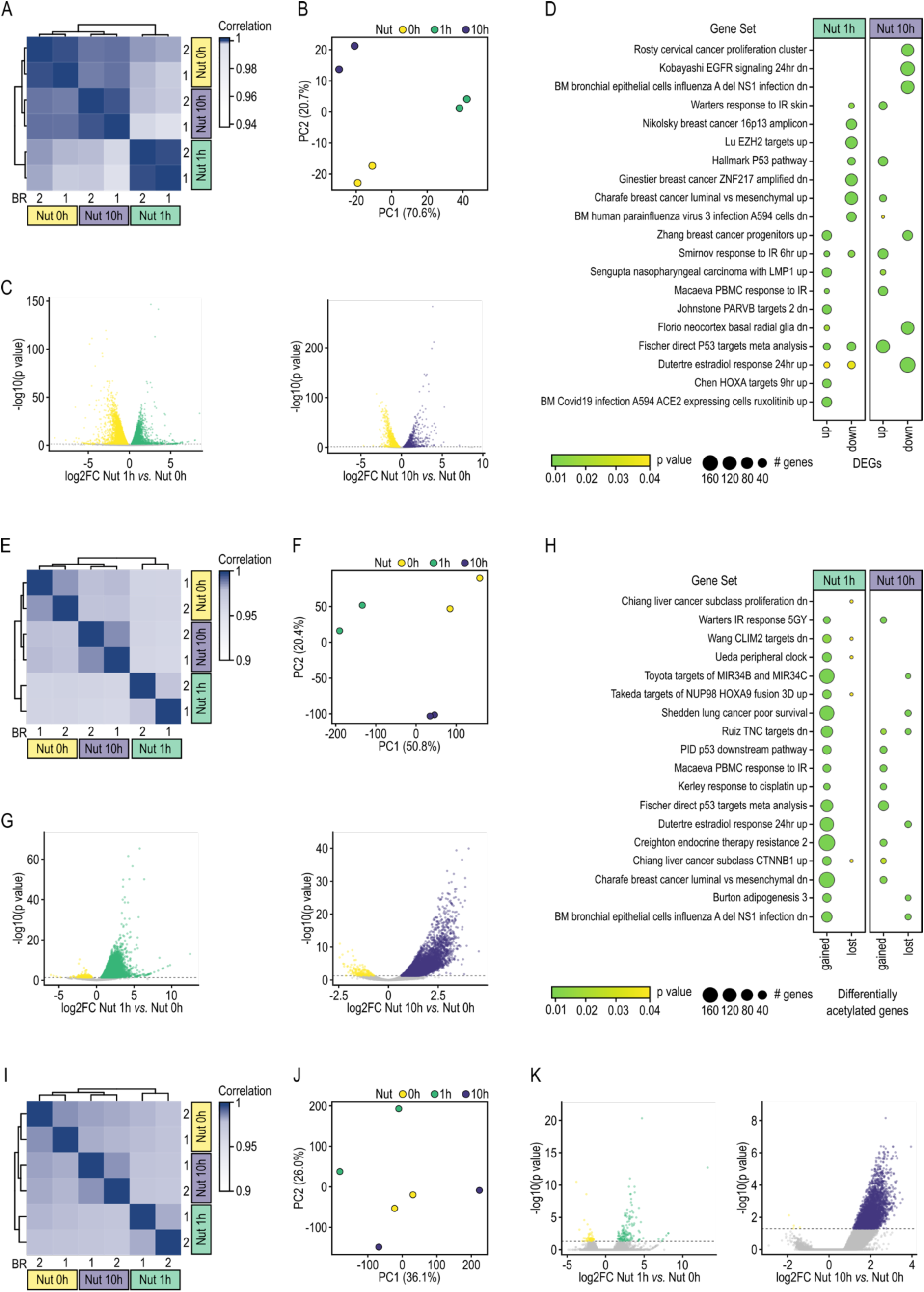
**A.** Pearson Correlation between biological replicates (1 and 2) of RNA-seq samples along p53 activation. Samples were clustered hierarchically inputting Euclidean distances to the “complete” agglomeration method. Nut 0h refers to control cells without p53 activation. **B.** Principal Component Analysis of RNA-seq biological replicates along p53 activation. Numbers in parentheses represent the percentage of variance explained by each principal component (PC). **C.** Differential expression analysis between 1 hour (Nut 1h) and 0 hour (Nut 0h) time points (left) and 10 hour (Nut 10h) and 0 hour (Nut 0h) time points (right). Differentially expressed genes are those with log2 fold change (log2FC) ≠ 0 and adjusted P value ≤ 0.05 (horizontal dotted line). Upregulated genes are in green or purple, downregulated genes are in yellow. **D.** Gene sets from the Molecular Signature Database (MSigDB) with significant enrichment (P-adj < 0.05) in one or more gene groups defined using differentially expressed genes (DEGS; upregulated as up; downregulated as down) at Nut 1h and Nut 10h time points compared to Nut 0h. Bubble size indicates the number of genes found in a gene set. p-values were calculated using a hypergeometric distribution test and adjusted for multiple comparison using a Benjamini-Hochberg (BH). **E, F, G** and **H.** As panel A, B, C and D respectively but for H3K27ac ChIP-seq data. **I, J** and **K.** As panel A, B and C respectively but for H3K27ac ChIP-seq data.

**Suppl. Fig. 3.**
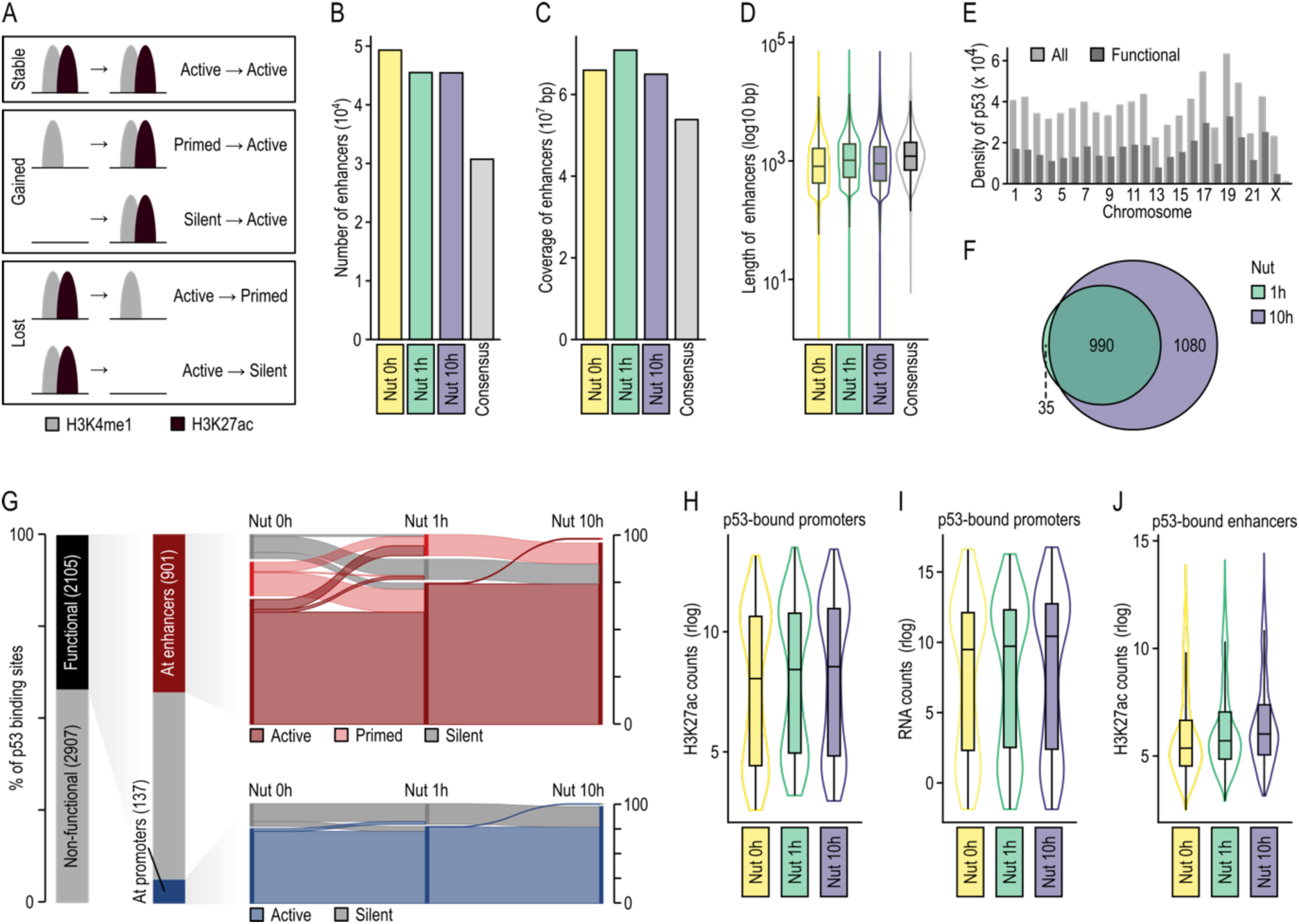
**A.** Schematic representation of our definition of enhancer dynamism. **B.** Number of enhancers defined at time points Nut 0h, Nut 1h and Nut 10h, as well as the consensus number of enhancers reached by intersecting the data of all three time points. Enhancers are defined as regions of the genome covered by H3K27ac and H3K4me1 peaks. Nut 0h refers to control cells without p53 activation. **C.** Genomic coverage of the enhancers defined at time points Nut 0h, Nut 1h and Nut 10h, as well as for the consensus set enhancers. **D.** Distribution of enhancer sizes defined at time points Nut 0h, Nut 1h and Nut 10h, as well as in the consensus set of enhancers. **E.** Number of all (light grey) and functional (dark grey) p53 binding sites per chromosome. Functional p53 binding sites are defined as p53 binding sites with an overlapping H3K27ac peak at either Nut 1h or Nut 10h time points. **F.** Overlap between functional p53 binding sites defined at Nut 1h and Nut 10h time points. **G.** Descriptive representation of the defined proportion of functional p53 binding sites (black) found at either promoters (blue) or enhancers (red) with varying states of activity over defined time points (Nut 0h, Nut 1h and Nut 10h). **H.** Distribution of normalized and rlog transformed H3K27ac counts at active promoter elements over the three time points (Nut 0h, Nut 1h and Nut 10h). **I.** Distribution of normalized and rlog transformed RNA-seq counts of genes with active promoter elements along p53 activation. **J.** Distribution of normalized and rlog transformed H3K27ac counts at active enhancers over the three time points (Nut 0h, Nut 1h and Nut 10h).

**Suppl. Fig. 4.**
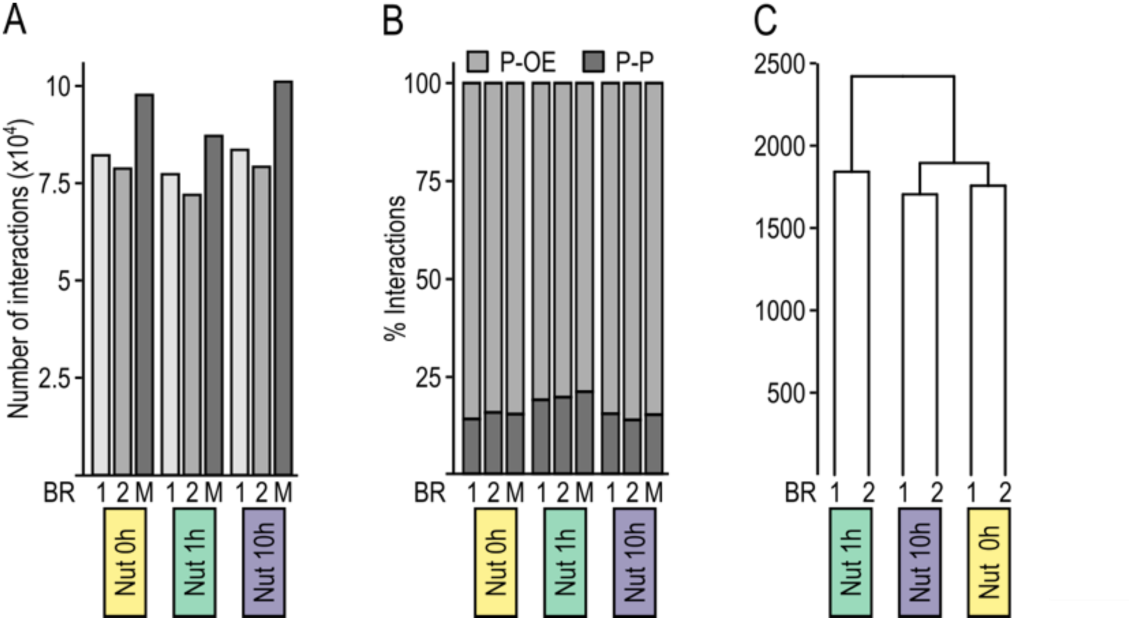
**A.** Total number of significant promoter interactions (CHICAGO score ≥ 5) of biological replicates (1 and 2) and merged samples (M) obtained by PCHi-C at Nut 0h, Nut 1h and Nut 10h time points. Nut 0h refers to control cells without p53 activation. **B.** Percentage of promoter-promoter (P-P) and promoter-other end (P-OE) significant interactions of biological replicates (1 and 2) and merged samples (M) obtained by PCHi-C along p53 activation. **C.** Hierarchical clustering of biological replicates (1 and 2) of promoter interactomes obtained by PCHi-C at Nut 0h, Nut 1h and Nut 10h time points. Euclidean distances were measured between samples, and the “complete” agglomeration method was used.

**Suppl. Fig. 5.**
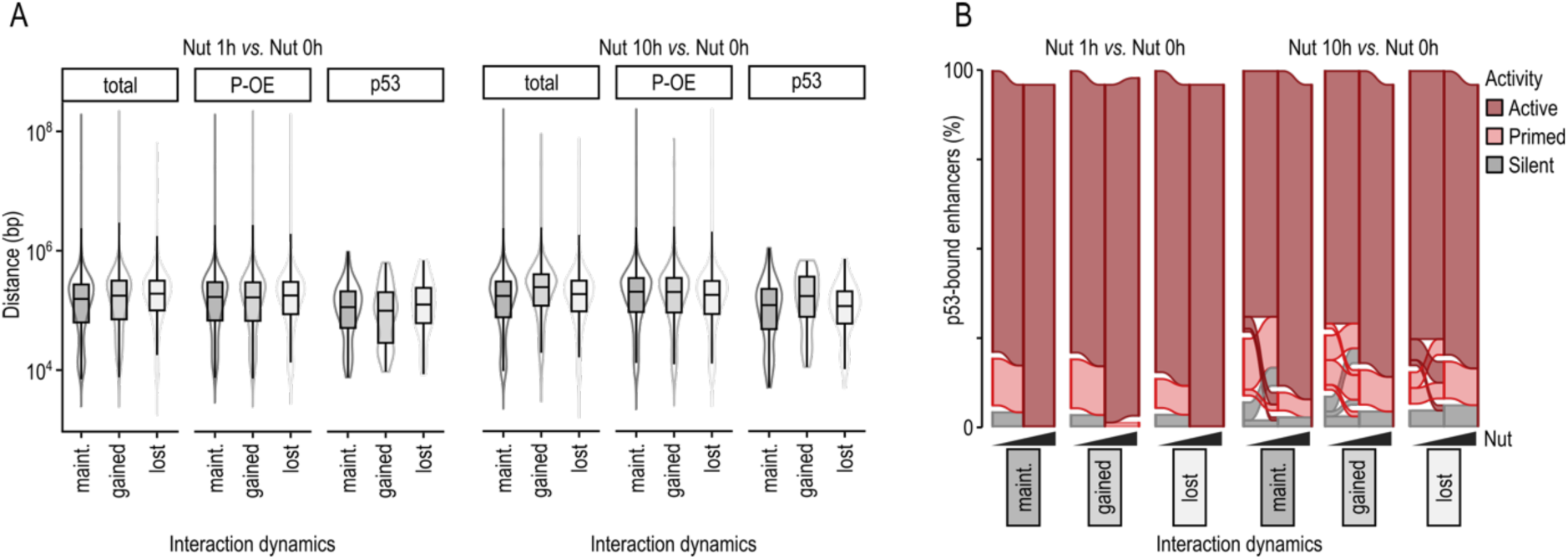
**A.** Distribution of distances of significant promoter interactions (CHiCAGO score ≥ 5) maintained, gained and lost between Nut 1h and Nut 0h time points (left) and Nut 10h and Nut 0h time points (right) respectively. The three interaction sets shown are comparing the complete interactomes (total); comparing interactions between a promoter and a non-promoter region (promoter with other-end or P-OE); and comparing interactomes with a p53 binding site present at either end (p53). Nut 0h refers to control cells prior to p53 activation. **B.** Sankey plot showing the activity dynamics of p53-bound enhancers after 1 hour (left) and 10 hours of p53 activation (right). Trends are divided considering only interactions maintained, gained or lost.

**Suppl. Fig. 6.**
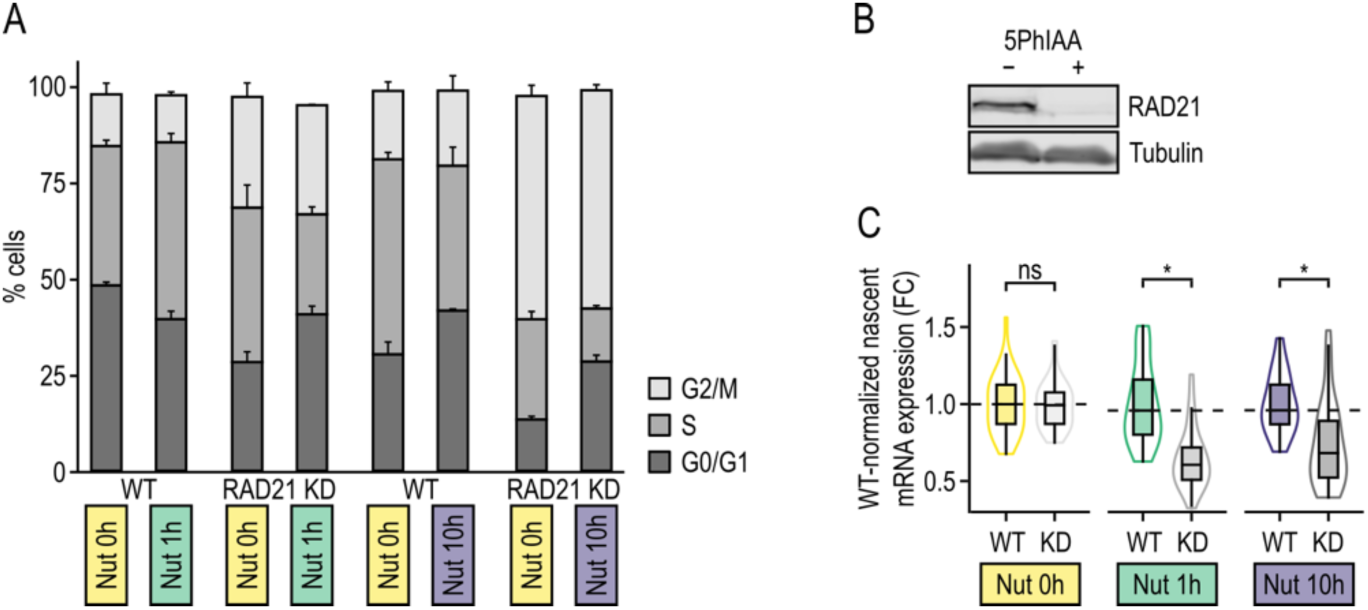
**A.** Representative distributions of cells in each cell cycle phase (G0–G1, S, G2–M) during Nutlin-3a treatment in the presence (WT) or absence of RAD21 (RAD21 KD). **B.** Western blot of RAD21 at with and without auxin treatment (*i.e.,* 5PhIAA), using Tubulin as control. **C.** Distribution of wild-type-normalized nascent mRNA expression levels (fold change) of distal target genes along p53 activation in wild type (WT) or RAD21 depleted (KD) conditions. A two-side Wilcoxon test was used to test whether expression levels differed significantly between conditions for each time point (star).

